# Orpinolide disrupts a leukemic dependency on cholesterol transport by inhibiting the oxysterol-binding protein OSBP

**DOI:** 10.1101/2023.03.15.532743

**Authors:** Marko Cigler, Hana Imrichova, Fabian Frommelt, Laura Depta, Andrea Rukavina, Chrysanthi Kagiou, J. Thomas Hannich, Cristina Mayor-Ruiz, Giulio Superti-Furga, Sonja Sievers, Luca Laraia, Herbert Waldmann, Georg E. Winter

## Abstract

Metabolic alterations in cancer precipitate in associated dependencies that can be therapeutically exploited. To meet this goal, natural product inspired small molecules can provide a resource of invaluable chemotypes. Here, we identify orpinolide, a synthetic withanolide analog with pronounced anti-leukemic properties via orthogonal chemical screening. Through multi-omics profiling and genome-scale CRISPR/Cas9 screens, we identify that orpinolide disrupts Golgi homeostasis via a mechanism that requires active phosphatidylinositol 4-phosphate (PI4P) signaling at the endoplasmic reticulum (ER)-Golgi membrane interface. Thermal proteome profiling and genetic validation studies reveal the oxysterol-binding protein OSBP as the direct and phenotypically relevant target of orpinolide. Collectively, these data reaffirm sterol transport as a therapeutically actionable dependency in leukemia and motivate ensuing translational investigation via the probe-like compound orpinolide.

## Introduction

Rewiring of metabolic networks contributes to the development, progression, and resistance acquisition of many cancer types, including leukemia, and is considered to be a general hallmark of cancer.^1, 2^ Patients with acute myeloid leukemia (AML) or acute lymphoblastic leukemia (ALL) display several metabolic changes, including alterations in lipid and cholesterol metabolism to enable the rapid proliferation of malignant cells, thus establishing motivation to exploit associated metabolic dependencies.^3, 4^ A suite of genome-scale genetic perturbation screens, for instance via CRISPR/Cas9 knockout screens, provide a rich catalogue of cell-autonomous, essential cancer cell functions, including dependencies on lipid or sterol metabolic pathways. However, redundancy and genetic buffering inherent to biological systems often requires simultaneous modulation of the activity of more than one protein, or modulation that goes beyond functional inhibition. These characteristics are often outside the reach of genetic perturbations but are attainable via small molecules.

Natural products (NPs) are an invaluable source of bioactive molecules with remarkable therapeutic potential. Approximately a third of all Food and Drug Administration (FDA) approved drugs in the last three decades originated from NPs or their derivatives.^5^ The evolutionary preconditioned and fine-tuned biological relevance of NPs is defined by the rich and diverse chemical space embedded in their structures.^6^ Yet, while advantageous, this structural complexity often encumbers the target identification and optimization of NP-derived bioactive compounds. To overcome these limitations, new strategies for identification and synthesis of simplified NP scaffolds are required.

The expansion of tangible chemical space can on the one side be achieved through biology-oriented synthesis (BIOS).^7^ BIOS simplifies NPs into core scaffolds with retained biological activities. These scaffolds in turn serve as starting points for the synthesis and further diversification of focused NP-inspired compound collections. On the other hand, fragmentation of NPs and their *de novo* combination allows the design of naturally non-occurring scaffolds, so-called pseudo-NPs.^8^ In both cases, fine-tuning of chemical space through simplification of complex parental NPs can yield compounds with improved or even unexpected biological activities, making them attractive starting points for drug discovery.

Withanolides are a class of plant-derived polyoxygenated steroidal lactones highly abundant in nature. Ever since the discovery and isolation of the archetypal withaferin A^9^, a plethora of different biological activities has been attributed to withanolides. Most notably, withaferin A and other derivatives have been shown to possess potent anti-inflammatory and anti-tumor effects by modulating cell signaling pathways such as NF-κB, MAPK and Wnt, or inhibiting the ubiquitin-proteasome system.^10–13^ It is thus not surprising that the withanolide scaffold has inspired the design of focused compound collections, prompting the identification of inhibitors of Hedgehog or Wnt signaling.^14, 15^ Noteworthy, withanolide D has been reported to induce apoptosis in leukemia cells through modulating ceramide production, while statin-mediated inhibition of the mevalonate pathway has recently been shown to be efficacious in early T cell progenitor ALL (ETP-ALL).^16, 17^ Collectively, these data establish a rationale to broadly assay the efficacy of withanolide-inspired small molecules in cellular models of AML and ALL.

Here, we couple leukemia cell line viability screens with morphological profiling of a focused withanolide-inspired compound collection and identify a cluster of structurally similar derivatives that show a pronounced anti-proliferative effect. This provided motivation to conduct unbiased, multi-omics target identification studies for the lead compound orpinolide (W7). Integrating quantitative proteomics and transcriptomics was indicative of functional impairment of Golgi homeostasis and cholesterol biosynthesis. To map cellular effectors that are required for the anti-proliferative consequences of orpinolide, we conducted a genome-scale CRISPR/Cas9 positive selection screen that revealed a requirement of active phosphatidylinositol 4-phosphate (PI4P) signaling at ER-Golgi membrane contact sites (MCSs). Using thermal proteome profiling (TPP), we ultimately uncover oxysterol-binding protein (OSBP) and its close ortholog ORP4 as direct targets of orpinolide over several other oxysterol-related protein (ORP) family members. Importantly, we verify that OSBP is the predominant metabolic dependency in leukemia cells, while ORP4 was largely dispensable. Altogether, through deciphering the cellular targets of withanolide-inspired orpinolide we reaffirm sterol transport at ER-Golgi MCSs as a potentially druggable vulnerability in leukemia cells.

## Results

### Phenotypic profiling of a withanolide-inspired compound collection

To systematically probe the effects of withanolides on leukemia cells, we selected a previously described compound collection based on the core steroid scaffold of type A withanolides (Extended Data Fig. 1).^15^ Generated by BIOS, the structural diversity within this focused library was predominantly introduced by derivatizing the δ-lactone side-chain while keeping the core sterol skeleton mostly unmodified. Previously studied in the context of cell signaling pathway modulators, several derivatives were shown to inhibit Wnt/β-catenin signaling in cell-based assays.^15^

The 52-membered compound collection was profiled in a cell line panel consisting of main lineages of leukemia: chronic myeloid (K562, KBM7), acute myeloid (MV4;11, OCIAML3) as well as pre-B cell (NALM6) and T cell acute lymphoblastic leukemia (Jurkat, LOUCY, MOLT4, P-12 Ichikawa). Following a 72 h treatment at 8 drug concentration points, we identified a cluster of compounds (W5-W11) that showed a pronounced cytotoxicity effect across all tested cell lines with the exception of K562 (Figure 1a, Extended Data Fig. 2 and Suppl. Table 1). Intriguingly, W5-W11 are all vinylogous urethane analogues derived from aliphatic (hetero)cyclic primary or secondary amines (Figure 1b), suggesting that the observed viability phenotype in leukemia requires certain structural/chemical features. This is further supported by the notion that, for example, the putative hydrolysis product of W5-W11 (W23) or the ring-fragmented product (W52) were not toxic to the tested cell lines (Extended Data Fig. 2).

**Figure 1.**
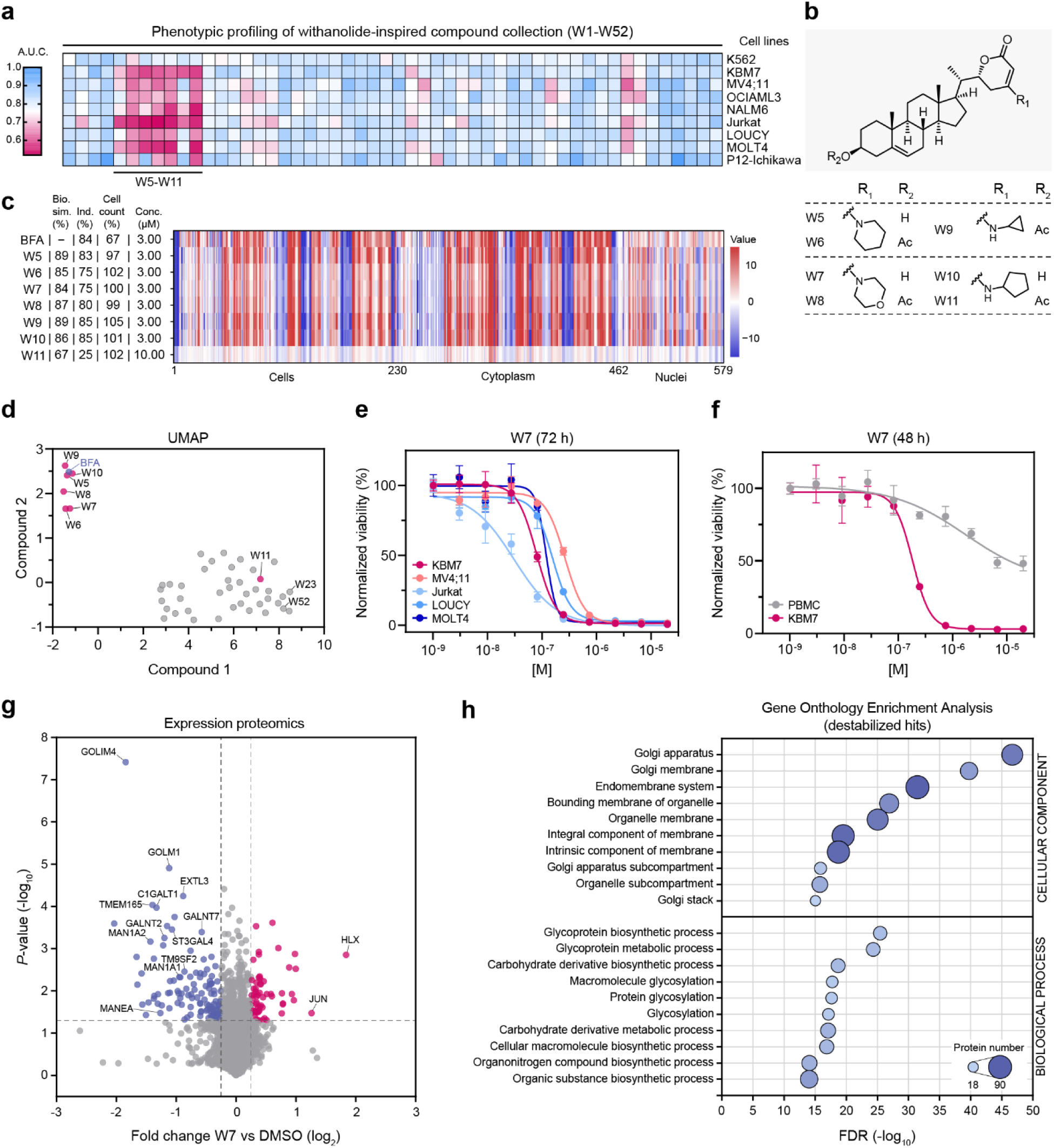
Orthogonal chemical profiling of a withanolide-inspired compound collection identifies W7 as disruptor of Golgi-related metabolic proteins. **(a)** Heatmap depicting results of the luminescence-based cell viability screening of a 52-membered withanolide-inspired compound collection in a panel of leukemia cells. Results are represented as relative area under the curve (A.U.C.) calculated from 8-concentration-point dose-response curves after 72 h drug treatment. **(b)** Chemical structures of seven derivatives (W5-W11) showing pronounced cytotoxicity in most of the tested cell lines. **(c)** Cell painting profiles of brefeldin A (BFA) and withanolides W5-W11 over 579 cellular features. Biosimilarity (Bio. sim.) values represent the similarity of phenotypic profiles while induction (Ind.) values represent the fraction of significantly changed cellular features. (d) UMAP analysis of the cell painting assay data highlighting the biological similarity of withanolides W5-W10 (3 μM, in fuchsia) to BFA (3 μM, in blue) over other withanolides (3 or 10 μM, in grey). The bioactivity fingerprint of the cytotoxic analogue W11 (10 μM, in fuchsia) differs from BFA. (e) Dose-resolved, normalized viability of the indicated leukemia cell lines after 72 h of W7 treatment (EC_50_ (KBM7) = 79.7 nM; EC_50_ (MV4;11) = 265.3 nM; EC_50_ (Jurkat) = 30.7 nM; EC_50_ (LOUCY) = 158.5 nM; EC_50_ (MOLT4) = 119.5 nM). Mean ± s.e.m.; n = 3 independent treatments. (f) Dose-resolved, normalized viability after 48 h treatment of blood-isolated peripheral blood mononuclear cells (PBMCs) or KBM7 with W7 (EC_50_ (KBM7) = 182.7 nM). Mean ± s.e.m.; n = 3 independent treatments. (g) Volcano plot of DMSO-normalized quantitative expression proteomics after 8 h treatment of KBM7 cells with W7 (1 μM). Proteins with higher or lower abundances in response to W7 treatment are marked in fuchsia or blue, respectively (adj. *P*-value < 0.05, log_2_ fold change greater than 0.25 or less than −0.25). (h) Gene Ontology (GO) enrichment analysis of destabilized expression proteomics hits (highlighted blue in (g)) for GO cellular components and GO biological processes. False discovery rate (FDR) (-log_10_-transformed) against the top 10 enriched terms sorted by adj. *P*-value is shown.

Next, we employed the cell painting assay (CPA)^18^ to characterize morphological phenotypes induced by W5-W11. To contextualize the obtained morphological profiles, we compared the bioactivity fingerprints of the tested withanolides to a set of reference compounds with known mode of action (MoA), looking for possible biological similarities between them. To our surprise, six out of seven cytotoxic analogues (W5-W10) showed a high similarity (> 80%) to the fungal metabolite brefeldin A (BFA) (Figure 1c). Notably, the morphological similarity was confined to the subset of withanolides that showed strong antileukemic properties (with the exception of W11), as visible in the UMAP analysis of recorded fingerprints (Figure 1d). This is particularly interesting when considering that BFA “glues” the interaction between the GDP-bound small GTPase Arf1 and its guanine nucleotide exchange factor ARNO.^19, 20^ Unable to release GDP, the stabilized “dead-end” conformation ultimately causes the disruption of the Golgi apparatus.^21, 22^ While the similarity to BFA does not imply that cytotoxic withanolides share the same molecular targets, it is indicative of an involvement of the Golgi in the anti-leukemic properties of W5-W10. To further elucidate this connection, we prioritized W7 and assessed its cellular efficacy in dose-ranging validation experiments (Figure 1e). W7 was particularly toxic in KBM7 and T-cell leukemia-derived cells with lower nanomolar EC50 values. Importantly, W7 did not substantially affect the viability of nonmalignant, peripheral blood mononuclear cells (PBMCs), thus highlighting a potential therapeutic window (Figure 1f). Of note, unlike other vinylogous urethane withanolides, W7 did not show any activity in cell-based Wnt pathway inhibition assays.^15^

### W7 treatment induces the destabilization of Golgi-related proteins and transcriptional repression of cholesterol biosynthesis

To understand and identify global proteome changes upon W7 treatment, we performed quantitative proteomics in KBM7 cells via the tandem mass tag (TMT) isobaric labeling approach. Out of over 7,800 quantified proteins, 175 displayed altered abundances (log2 fold change greater than 0.25 or less than −0.25) after 8 h treatment with W7 (Suppl. Table 2). Strikingly, a substantial number of proteins (116, 66%; adj. *P*-value < 0.05) was strongly destabilized (Figure 1g). Functional analysis of destabilized hits revealed a high enrichment of Golgi-localized transmembrane proteins (e.g. GOLIM4, GLG1, TM9SF1-4) as well as proteins involved in glycoprotein biosynthesis (e.g. GALNT2/7, MAN1A1/2, C1GALT1) (Figure 1h). To increase the confidence in the observed phenotype, we investigated if destabilized proteins are known to engage in protein-protein interactions or participate in shared protein complexes. To this end, we employed the open-access BioGRID database^23^ to create a protein-protein interaction (PPI) network between hits whose abundances were significantly altered upon W7 treatment. For more than half of the perturbed proteins (97 of 175 or 55%) direct interactions were reported in the literature which represents significantly more shared edges than a random sampling of proteins would result in (Extended Data Fig. 3a-3c, Suppl. Table 2). Mapping of the subcellular localization of the proteins within the network illustrates the overall impact on the Golgi proteins as a phenotypic response to W7 treatment. Moreover, we validated the W7-induced time-dependent destabilization of the endosomal cycling protein and trafficking chaperone Golgi integral membrane protein 4 (GOLIM4) (Extended Data Fig. 3d). Collectively, our data indicates that W7 disrupts the integrity of the Golgi apparatus, further supporting morphologic features detected via CPA. The molecular mechanism behind global destabilization of related metabolic (especially glycosylating) and trafficking proteins, however, remained elusive.

To further explore the mechanism of action of W7, we set out to examine the global transcriptional effects upon W7 treatment. We performed mRNA-sequencing of vehicle (DMSO) or compound treated KBM7 cells (Figure 2a, Suppl. Table 3). While we observed an upregulation of several genes involved in the JAK-STAT pathway (e.g. STAT4, cytokine receptors such as IL-2), detailed analysis of the differential gene expression profile of W7 revealed distinct changes in a subset of important metabolic genes. More precisely, the transcriptional response to W7 treatment included a significant downregulation of *de novo* cholesterol biosynthesis genes such as *HMGCS1*, *LSS*, *DHCR7* and *SQLE*, which encodes for one of the two rate-limiting enzymes in this process (Figure 2a-2b). Moreover, key regulators of cholesterol metabolic processes *SREBF2* and *INSIG1* were downregulated as well. It is worth noting that similar repression of cholesterol production can be induced by oxysterols such as 25- and 27-hydroxycholesterol.^24^ These endogenous cholesterol oxidation products suppress the transcriptional response of SREBP2 by hindering its trafficking to the Golgi and thereby preventing its proteolytic processing and maturation.^25, 26^

**Figure 2.**
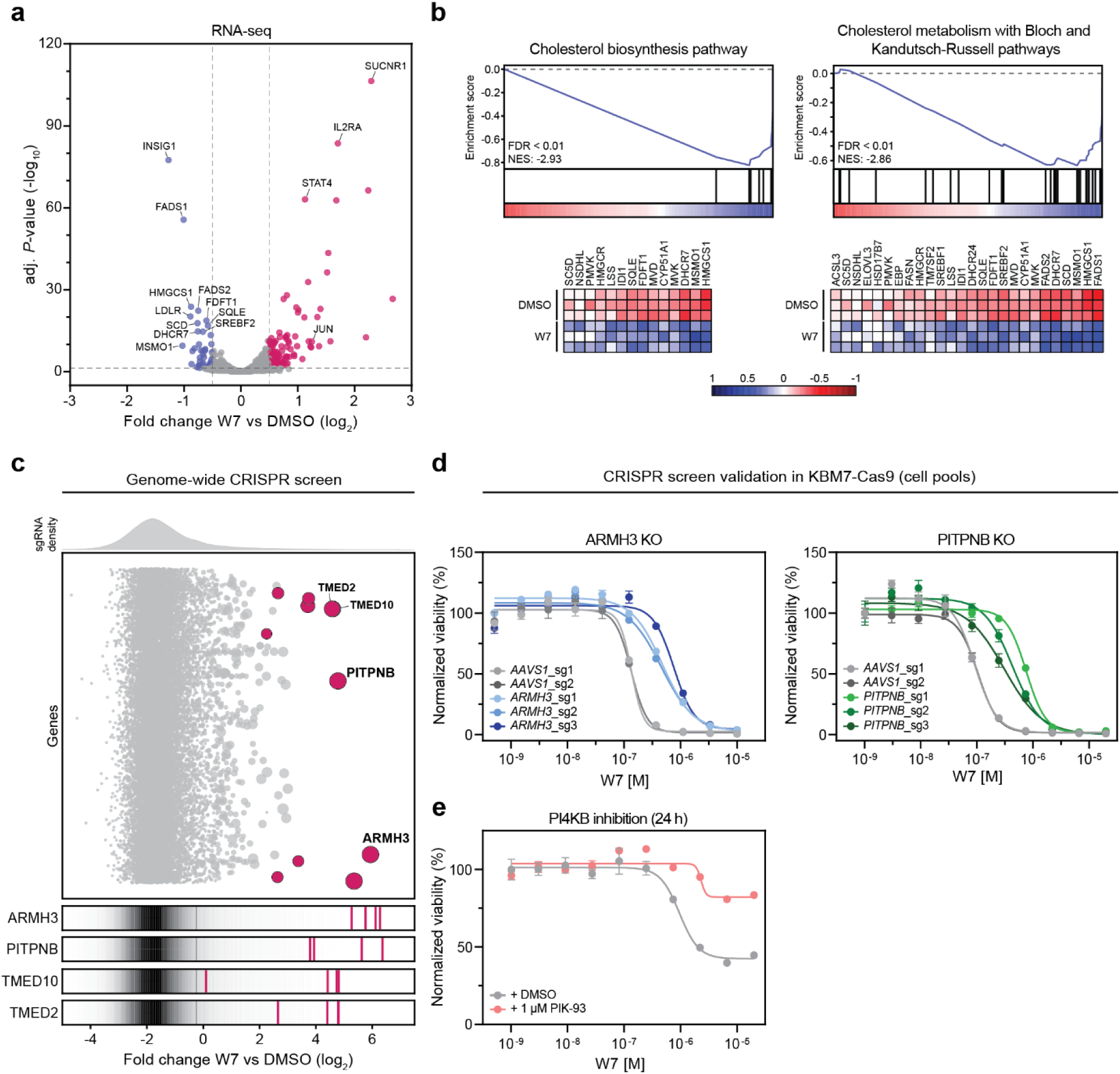
W7 inhibits cholesterol biosynthesis and functionally depends on PI4P signaling. **(a)** Changes in gene expression after 6 h treatment of KBM7 cells with W7 (485 nM). Downregulated and upregulated genes (adj. *P*-value < 0.05, log_2_ fold change greater than 0.5 or less than −0.5) are marked in blue or fuchsia, respectively. **(b)** Gene set enrichment analysis (GSEA) of top two WikiPathway gene sets from MSigDB significantly enriched in the differential gene expression profile of W7 against DMSO. Normalized enrichment scores (NES) and false discovery rate (FDR) are shown. VST-normalized median-centered gene expression of the leading edge subset of genes are depicted in the corresponding heatmaps. Genes are ordered based on the metascore. **(c)** Genome-wide CRISPR/Cas9 knockout screen in W7-treated KBM7 cells constitutively expressing Cas9 (KBM7-Cas9). The bubble plot displays median sgRNA enrichment upon W7 treatment over DMSO while the bubble size represents significance. Only genes with FDR ≤ 0.05 are highlighted. Enrichment of sgRNAs targeting selected hits in comparison to the distribution of all sgRNAs are highlighted separately. **(d)** Dose-resolved, normalized viability after 72 h treatment of KBM7-Cas9 cells expressing sgRNAs targeting *ARMH3* (left, in blue), *PITPNB* (right, in green) or *AAVS1* locus (grey) with W7. Mean ± s.e.m.; n = 3 independent treatments. **(e)** Dose-resolved, normalized viability of KBM7 cells with W7 in the presence (red) or absence (grey) of PI4KB inhibitor PIK-93 (1 μM). The cells were co-treated for 24 h. Mean ± s.e.m.; n = 3 independent treatments.

Collectively, multi-omics profiling reveals that W7 treatment disrupts cholesterol biosynthesis and affects the cellular abundance of an array of Golgi-associated proteins, indicative of a functional impairment of Golgi homeostasis.

### Cellular efficacy of W7 functionally depends on PI4P signaling between the ER and the Golgi

Next, we wanted to investigate whether the observed impact on Golgi morphology is cause or consequence of the cellular efficacy of W7. To determine the cellular effectors required for the anti-proliferative potential of W7 in leukemia, we performed a genome-wide CRISPR/Cas9 knockout screen in KBM7 cells.^27^ After puromycin selection, the transduced cell pools were continuously treated with W7 or vehicle (DMSO) over 20 days, thus enabling the enrichment of genes functionally required for the anti-proliferative response to the compound (Extended Data Fig. 4a, Suppl. Table 4). Strikingly, top hits included *ARMH3* (until recently C10orf76) and *PITPNB* (Figure 2c). Both genes constitute the molecular machinery required for maintaining phosphatidylinositol 4-phosphate (PI4P) signaling between the ER and other membrane organelles like the trans-Golgi network (TGN). While phosphatidylinositol (PI) transfer protein beta (PITPNB) mediates the ER-to-TGN transport of PI,^28, 29^ Armadillo-like helical domain-containing protein 3 (ARMH3) associates with PI 4-kinase beta (PI4KB) and plays an important role in PI4KB-driven PI4P generation at the Golgi.^30, 31^ In addition, sgRNAs targeting the components of membrane contact sites (MCSs) involved in lipid transport like transmembrane emp24 domain containing protein (TMED) 2 and 10^32^ were also enriched (Figure 2c). Targeted CRISPR-induced inactivation of *PITPNB* and *ARMH3* in KBM7 cell pools confirmed their requirement for W7 efficacy (Figure 2d). Similarly, chemical inhibition of PI4KB with PIK-93 rescued the W7-induced cytotoxicity in KBM7 cells (Figure 2e), corroborating the importance of active PI4P signaling at MCSs and supporting the CRISPR screening data. Apart from its active role in regulating membrane trafficking, the intrinsic gradient of PI4P (high within the TGN and low within the ER) also drives the non-vesicular transport of lipids, especially cholesterol, at MCSs. Several endogenous metabolites such as oxysterols are involved in PI metabolism but also contribute to the regulation of cholesterol homeostasis. Given that W7 induced transcriptional repression of cholesterol biosynthesis (Figure 2a-2b), we wanted to distinguish it from oxysterols like 25-hydroxycholesterol (25-OHC). Therefore, we probed the cellular efficacy of 25-OHC in *PITPNB* and *ARMH3* knockout background. In both cases, depletion of PITPNB and ARMH3 did not affect 25-OHC cytotoxicity in KBM7 cells (Extended Data Fig. 4b).

Taken together, our findings show that the cellular efficacy of W7 functionally depends on PI4P signaling between the ER and the Golgi. Given the essential role of PI4P in maintaining Golgi function, we reason that the observed impact on Golgi homeostasis is causative for the leukemic vulnerability to W7. Hence, we anticipated that the direct protein target(s) of W7 would be involved in Golgi-associated processes.

### Orpinolide (W7) targets the oxysterol-binding protein OSBP

The functional requirement of PI4P signaling led us to surmise that W7 might interfere with the sterol transport at ER-Golgi contact sites. To identify cellular targets of W7, we performed thermal proteome profiling (TPP) of KBM7 cells. Using multiplexed quantitative proteomics, TPP allows the assessment of proteome-wide changes in protein stability upon interactions with small molecules.^33^ Importantly, the distinct changes in the melting profiles of proteins are indicative of putative drug targets. In brief, after initial treatment with vehicle (DMSO) or W7, intact KBM7 cells were subjected to thermal destabilization with an 8-temperature gradient (37-62 °C). Following mild cell lysis, the samples were subjected to standard quantitative proteomics workflow via TMT isobaric labeling (Extended Data Fig. 5a, Suppl. Table 5). Subsequent nonparametric analysis of response curves (NPARC)^34^ revealed 4037 melting profiles, out of which 113 were significantly altered (adj. *P*-value < 0.01) (Extended Data Fig. 5b-5d, Suppl. Table 5). These included both stabilization (26) and destabilization (5) profiles with a Δ*T*m exceeding 1 °C (Figure 3a and Extended Data Fig. 5d). Considering the information embedded in the experimental data acquired hitherto, we overlaid the 31 discrete hits with the TGN-localized (UniProtKB subcellular localization SL-0266) and sterol-binding (GO:0032934) proteins. This intersection revealed oxysterol-binding protein (OSBP) as the putative target of W7 (Figure 3b, Extended Data Fig. 5e). As a major lipid transporter at ER-Golgi contact sites, OSBP utilizes the metabolic energy of PI4P to exchange cholesterol as well as PI4P between the membranes.^35, 36^ We confirmed drug binding to OSBP through two versions of cellular thermal shift assay (CETSA): via immunoblotting of endogenous OSBP in KBM7 cells (Figure 3c) as well as the split nanoluciferase (HiBiT-LgBiT) system in KBM7 cells overexpressing HiBiT-tagged OSBP (Figure 3d). With the latter approach we furthermore show the dependency of OSBP stabilization on W7 concentration (Extended Data Fig. 6a). TPP and CETSA experiments thus point to OSBP as direct protein target of W7, prompting us to term the compound *orpinolide* henceforth. To confirm target engagement in an *in vitro* setup, we conducted fluorescent polarization (FP) assays with recombinant GST-tagged sterol-binding domain of OSBP (OSBP-related domain or ORD), which indicated that orpinolide engages OSBP in the nanomolar range (Figure 3e). Employing a suite of additional FP assays^37, 38^, we could show that orpinolide binding is confined to OSBP over other sterol transporters such as the members of Aster and STARD families (Extended Data Fig. 6b). OSBP, however, is the prototypical member of the OSBP-related protein (ORP) family: dual lipid transporters at MCSs characterized by possessing conserved ORD domains.^39^ In order to profile orpinolide specificity within this protein family, we overexpressed several different ORPs (ORP1, ORP2, full-length variant of ORP4 or ORP4L, ORP9 and ORP11) as HiBiT-fusions (Extended Data Fig. 6c) and assessed orpinolide binding via HiBiT-CETSA in intact KBM7 cells. We particularly focused on ORPs known for binding cholesterol (based on reported *in vitro* or *in vivo* data) and/or Golgi localization.^40^ We observed that orpinolide stabilized the closely related ortholog ORP4L, albeit to a lower degree (Figure 3f). In contrast, none of the other tested ORPs were thermally stabilized in the presence of the compound (Extended Data Fig. 6d). Of note, finding OSBP (and ORP4L) as putative target(s) of orpinolide prompted us to reexamine the expression proteomics dataset (Figure 1g). While OSBP and other ORPs were detected in the experiment, their relative abundances were unchanged upon compound treatment (Suppl. Table 2) which we could also confirm in case of OSBP via immunoblotting (Extended Data Fig. 6e). Collectively, these data establish OSBP/ORP4L as cellular targets of orpinolide.

**Figure 3.**
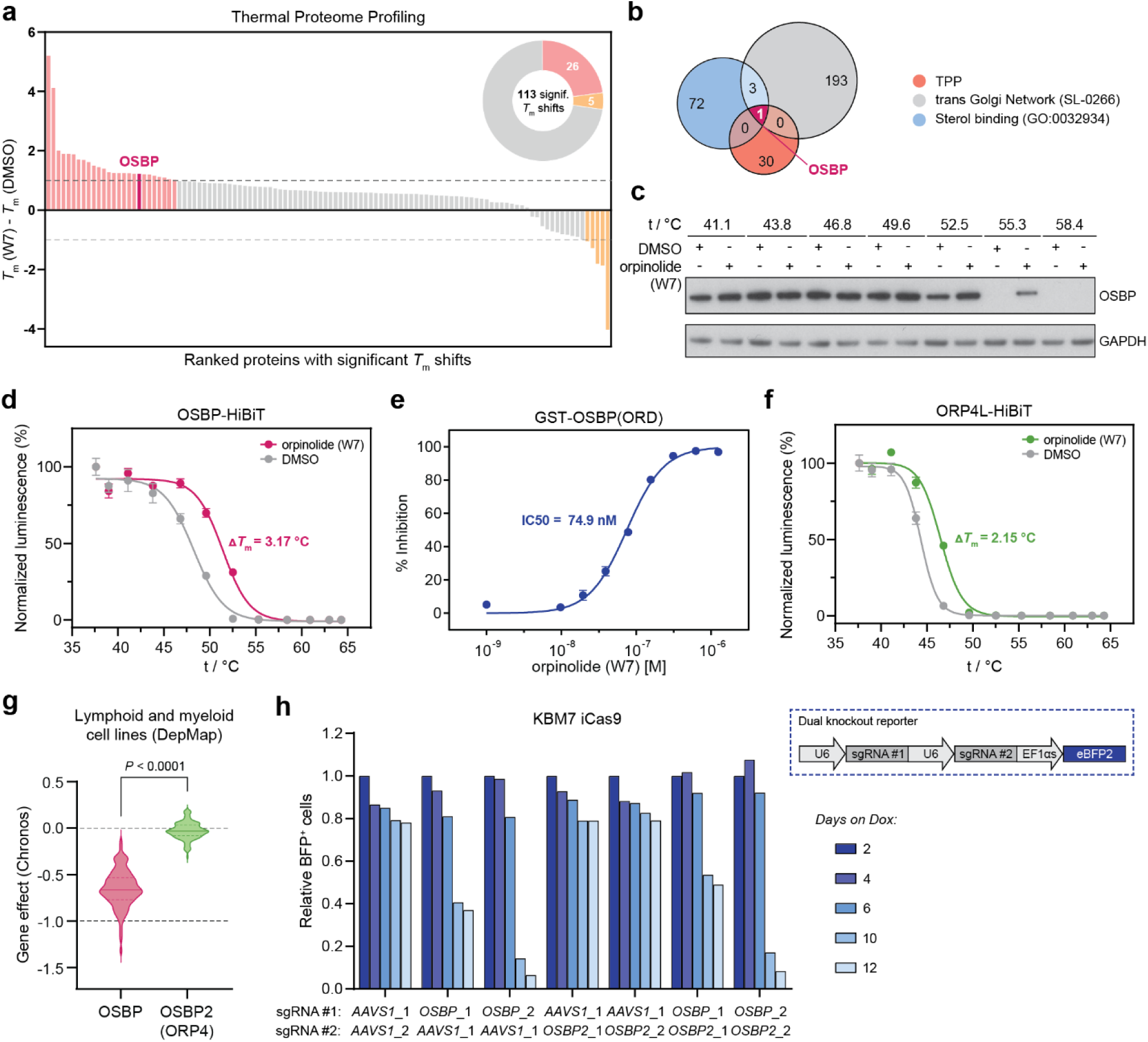
OSBP is the direct and phenotypically relevant target of orpinolide (W7) **(a)** Waterfall plot depicting the results of thermal proteome profiling (TPP). Out of 113 significantly altered melting profiles, 31 discrete hits show a Δ*T*m exceeding 1 °C (colored in red and orange). **(b)** Overlay of the 31 hits from TPP with TGN localized (UniProtKB subcellular localization SL-0266) and sterol-binding proteins (GO:0032934) reveals OSBP as the putative target of orpinolide (W7). **(c)** Orpinolide-induced thermal stabilization of endogenous OSBP in KBM7 cells. Intact cells were treated for 2 h with 5 μM orpinolide before performing the CETSA. Representative images of n = 2 independent measurements. **(d)** Orpinolide-induced thermal stabilization of HiBiT-tagged OSBP in KBM7 cells stably expressing the fusion. Intact cells were treated for 4 h with 1 μM orpinolide before performing the CETSA. Mean ± s.e.m.; n = 3 technical replicates; representative of n = 3 independent experiments is shown. **(e)** Fluorescence polarization competition assay measuring the displacement of 22-NDB-cholesterol from purified GST-tagged ORD of OSBP upon orpinolide treatment. Mean ± s.e.m.; n = 3 technical replicates; representative of n = 3 independent experiments is shown. **(f)** Orpinolide-induced thermal stabilization of HiBiT-tagged ORP4L in KBM7 cells stably expressing the fusion. Intact cells were treated for 4 h with 1 μM orpinolide before performing the CETSA. Mean ± s.e.m.; n = 3 technical replicates; representative of n = 2 independent experiments is shown. **(g)** Distribution of OSBP and ORP4 (*OSBP2*) deletion effects across myeloid and lymphoid cancer cell lines from the DepMap 22Q4 CRISPR data. **(h)** CRISPR-based dropout experiment in Dox-inducible Cas9 KBM7 (KBM7 iCas9) cells transduced with a BFP-based dual knockout reporter. The reporter plasmid carries a combination of sgRNAs targeting *OSBP*, *OSBP2* or *AAVS1* locus to yield single or dual *OSBP/OSBP2* knockout. The bar plots depict relative BFP_+_ cells over 12-day Dox treatment, normalized to day 2 for each individual reporter system.

Previous studies have implicated the dysregulation and/or aberrant expression of various members of the ORP family with the development of cancer.^41, 42^ OSBP and ORP4L have, for example, been revealed as targets of several natural products displaying anti-proliferative effects in diverse cancer cell lines.^43^ Moreover, the proliferation and survival of immortalized and cancerous cell lines, especially T cell-derived lymphoblastic leukemia, seemingly requires the expression of ORP4.^44, 45^ We thus wanted to investigate if functional interference with OSBP, ORP4 or both recapitulates the orpinolide-induced vulnerability phenotype of leukemia cells. To that goal, we turned again towards functional genomics. Closer inspection of the publicly available genome-wide CRISPR/Cas9 knockout screening data across 1078 cell lines unveiled ORP4-encoding *OSBP2* as a non-essential gene. In contrast, a distinct dependency on *OSBP* can be observed in multiple cell lines (Extended Data Fig. 6f), including ones originating from myeloid or lymphoid lineages (Figure 3g).^46^ To evaluate these pooled screening results in an arrayed format in the context of leukemia, we conducted a CRISPR-based dropout screen in KBM7 and Jurkat cells by inducing either single or dual *OSBP*/*OSBP2* knockout(s). By monitoring the decrease of the sgRNA (eBFP2-) positive cells over time, our results suggest that the survival of KBM7 cells solely depends on *OSBP* without significant contribution of *OSBP2* (Figure 3h). A similar effect of OSBP inactivation was observed in Jurkat cells (Extended Data Fig. 6g).

In summary, through various multi-omics approaches we uncover that orpinolide interferes with the sterol transport at ER-Golgi MCSs by selectively targeting OSBP and ORP4L over other ORPs. Importantly, we disentangle the cellular efficacy of orpinolide by showing that it predominantly emanates from targeting OSBP, reaffirming it as a putative therapeutic target in the context of leukemia.

## Discussion

Here, we assessed the anti-leukemic properties of a focused compound collection constructed based on the core withanolide scaffold and identified orpinolide amongst several vinylogous urethane analogs with pronounced anti-proliferative effects. To dissect its mode of action and identify its cellular target(s), we integrated quantitative proteomics, transcriptomics and functional genomics screens and revealed that orpinolide-treatment leads to functional impairment of Golgi homeostasis by targeting OSBP. Serving as a lipid transporter at ER-Golgi MCSs, OSBP utilizes the metabolic energy of PI4P to drive the transport of ER-produced cholesterol to the Golgi. At the same time, it helps maintain the intrinsic PI4P gradient between membranes by transferring PI4P back to the ER where it is rapidly dephosphorylated.^35, 36^ Hence, OSBP is an essential component of non-vesicular lipid transport at MCSs. Upon profiling the specificity of orpinolide against several other members of the ORP family, we identified ORP4 as its additional target. Like OSBP, ORP4 is involved in the sterol/PI4P counter transport.^44^

To the best of our knowledge, orpinolide represents a first withanolide-derived modulator of OSBP/ORP4 activity. OSBP/ORP4 are also bound by oxysterols, including 25-OHC. However, our results clearly differentiate the effect of orpinolide from 25-OHC where the functional dependence of orpinolide on active PIP4 signaling was not recapitulated by 25-OHC. The promiscuity of 25-OHC, which interacts with a plethora of sterol-sensing proteins^47, 48^ could serve as an explanation for this difference, but definitive elucidation will require further investigation. Along similar lines, several bioactive NPs have previously been reported as OSBP/ORP4 modulators and have collectively been referred to as “ORPphilins”.^43^ This includes compounds such as OSW-1 and schweinfurthin A. Noteworthy, their molecular mechanism of OSBP modulation appears to be highly diverse, not only in terms of binding potency and promiscuity, but also with respect to the mechanistic consequences following target engagement. For instance, OSW-1 treatment induced proteasome-dependent degradation of OSBP via an entirely elusive mechanism. In contrast, orpinolide has no consequence on OSBP stability or abundance. The specific and individual molecular consequences prompted by OSBP/ORP4 target engagement will hence require further elucidation. In addition, the synthesis of these natural products is time- and resource intensive, whereas orpinolide is accessible by means of a short and efficient synthesis sequence.^15^

Several studies have addressed the function of ORP4 in leukemia. Previous work has implied it as essential driver of T-ALL proliferation and found it to be overexpressed in malignant compared to benign T cells.^45^ Akin to OSBP, ORP4 is also thought to orchestrate the transport of PI4P between the ER and the plasma membrane, thereby supporting hyperactivity of the PI3K/AKT pathway and leukemogenesis of T cells.^49^ Initially, these observations provided an intriguing hypothesis for the observed anti-leukemic efficacy of orpinolide as a selective, dual inhibitor of OSBP/ORP4. Arrayed gene disruption experiments of OSBP and ORP4, either individually or in combination, however, clearly indicate that OSBP represents the predominant dependency. This is in agreement with genome-scale CRISPR/Cas9 screening data that is publicly available in DepMap.^46^ Our data implies that, even when assessed in combination with OSBP knockout, ORP4 disruption prompts no measurable fitness defect. OSBP-mediated cholesterol transport to lysosomes has also been implicated in sustaining aberrant mTORC1 signaling, particularly in disease models of Niemann-Pick disease, which is caused by loss of the lysosomal cholesterol transporter NPC1.^50^ The relevance of altered mTORC1 for the anti-leukemic properties of orpinolide will be the subject of future research.

Collectively, the presented data establishes orpinolide as a probe-like small molecule that can be employed to investigate cholesterol transport by OSBP and ORP4. Our data also reaffirms sterol transport at MCSs as a druggable metabolic dependency in leukemia cells that will motivate future translational efforts.

## Author contributions

M.C., H.W. and G.E.W. conceptualized this study. M.C. designed and conducted the experiments with help from C.K. and C.M.-R. H.I. analyzed and visualized RNAseq and CRISPR screening data. M.C. and A.R. performed the TPP experiment while F.F. analyzed and visualized the TPP data. F.F. performed the protein interaction analysis. L.D. developed and performed the recombinant binding assays under supervision of L.L. S.S. performed the cell painting assay. J.T.H. supervised the proteomics experiments. G.S.F. supervised TPP analyses. M.C. generated figures with input from all authors. M.C. and G.E.W. wrote the manuscript with input from all authors.

## Acknowledgements

We are grateful to all the members of the Winter lab, in particular Natalie Scholes, Alexander Hanzl and Vincenth Brennsteiner for helpful discussions and editorial contributions. We thank Anna Koren, Stefan Kubicek and the CeMM Molecular Discovery Platform for their assistance with the profiling of the withanolide-inspired compound collection; and the CeMM Biomedical Sequencing Facility for NGS sample processing, sequencing, and data curation. We moreover thank Johannes Zuber at the Research Institute of Molecular Pathology for sharing iCas9 cell lines and plasmids.

## Funding

CeMM, the Winter lab and the Superti-Furga lab are supported by the Austrian Academy of Sciences. The Winter lab is further supported by funding from the European Research Council (ERC) under the European Union’s Horizon 2020 research and innovation program (grant agreement 851478), as well as by funding from the Austrian Science Fund (FWF, projects P32125, P31690 and P7909). The Laraia Lab was supported by funding from the Novo Nordisk Foundation (NNF21OC0067188) and the Independent Research Fund Denmark (9041-00248B).

## Competing interests

G.E.W. and G.S.F. are scientific founders and shareholders of Proxygen and Solgate. The Winter and Superti-Furga labs received research funding from Pfizer. C.M-R. is part of the SAB of Nostrum Biodiscovery. The C.M.-R. lab receives research funding from Aelin Therapeutix and Almirall.

## Extended Data

**Extended Data Fig. 1.**
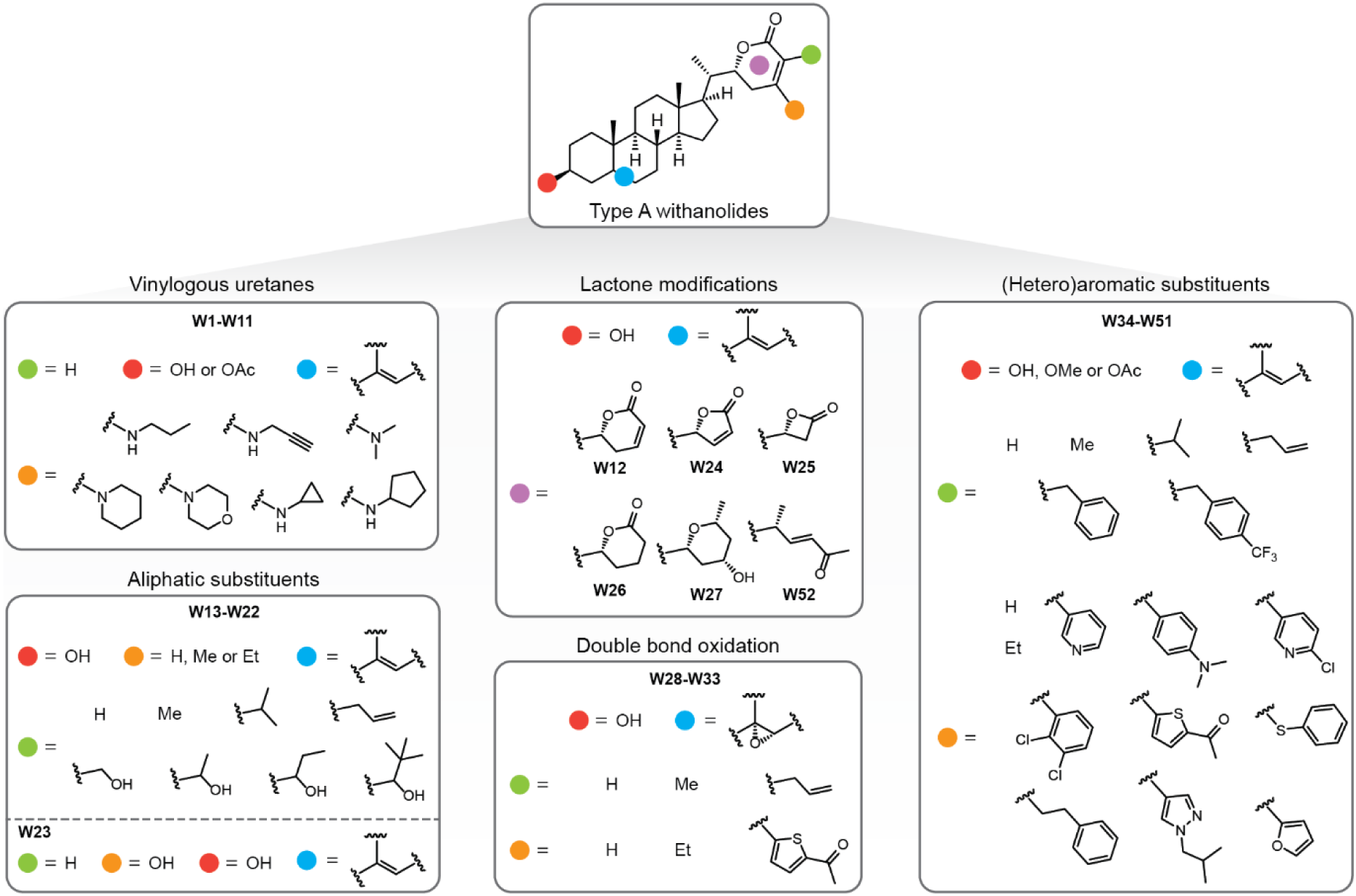
Withanolide-inspired compound collection. Structural overview of the 52-membered compound library based on the type A withanolide scaffold. SMILES representations of the compound collection are available in the Suppl. Table 1.

**Extended Data Fig. 2.**
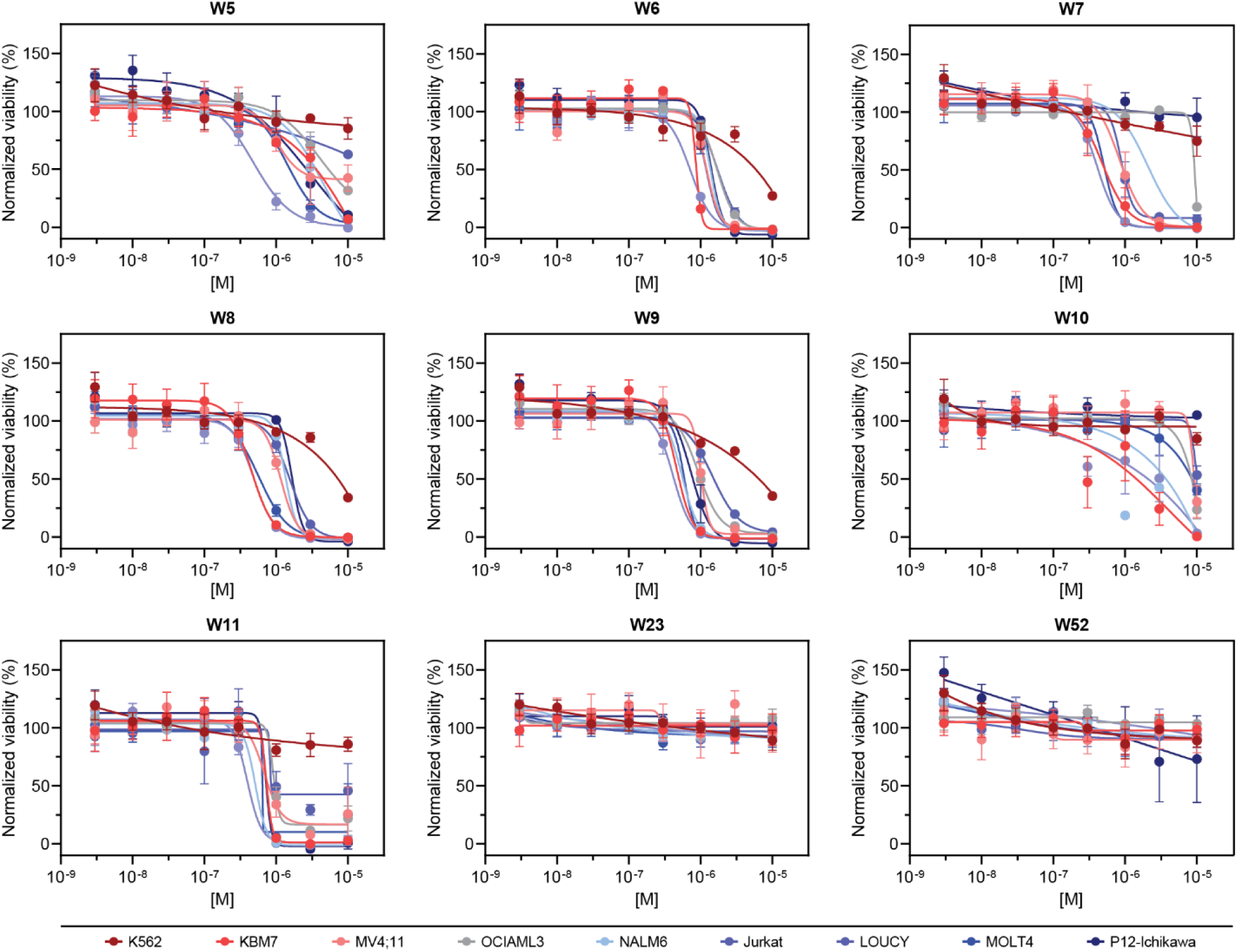
Phenotypic profiling of the withanolide-inspired compound collection in leukemia cells. Representative dose response curves for seven cytotoxic vinylogous urethans (W5-W11) as well as their putative hydrolysis product (W23) and the ring-fragmented product (W52). Mean ± s.e.m.; n = 3 independent treatments.

**Extended Data Fig. 3.**
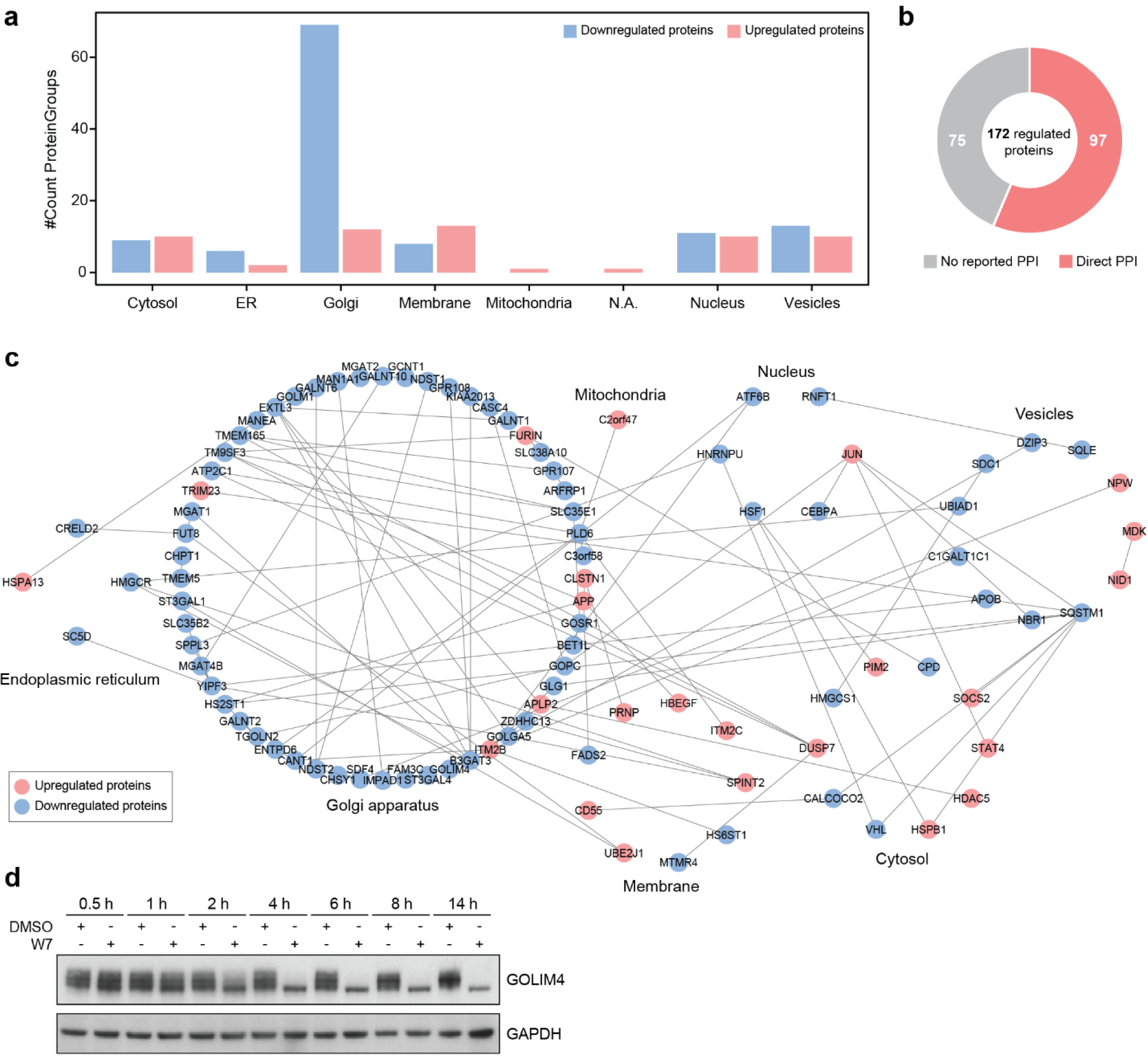
W7 induces global destabilization of Golgi-related proteins. **(a)** Subcellular localization distribution of protein hits with significantly altered abundances. Proteins were categorized based on the information from The Human Protein Atlas and UniProtKB. **(b)** Distribution of direct PPIs reported in the BioGRID database among the 172 regulated proteins reported in the BioGRID database. **(c)** Protein-protein interaction (PPI) network of proteins whose abundances were significantly altered upon W7 treatment. The physical interactions were derived from the BioGRID interaction database. **(d)** Time-dependent GOLIM4 destabilization induced by W7 treatment (1 μM) in KBM7 cells. Representative images of n = 2 independent measurements.

**Extended Data Fig. 4.**
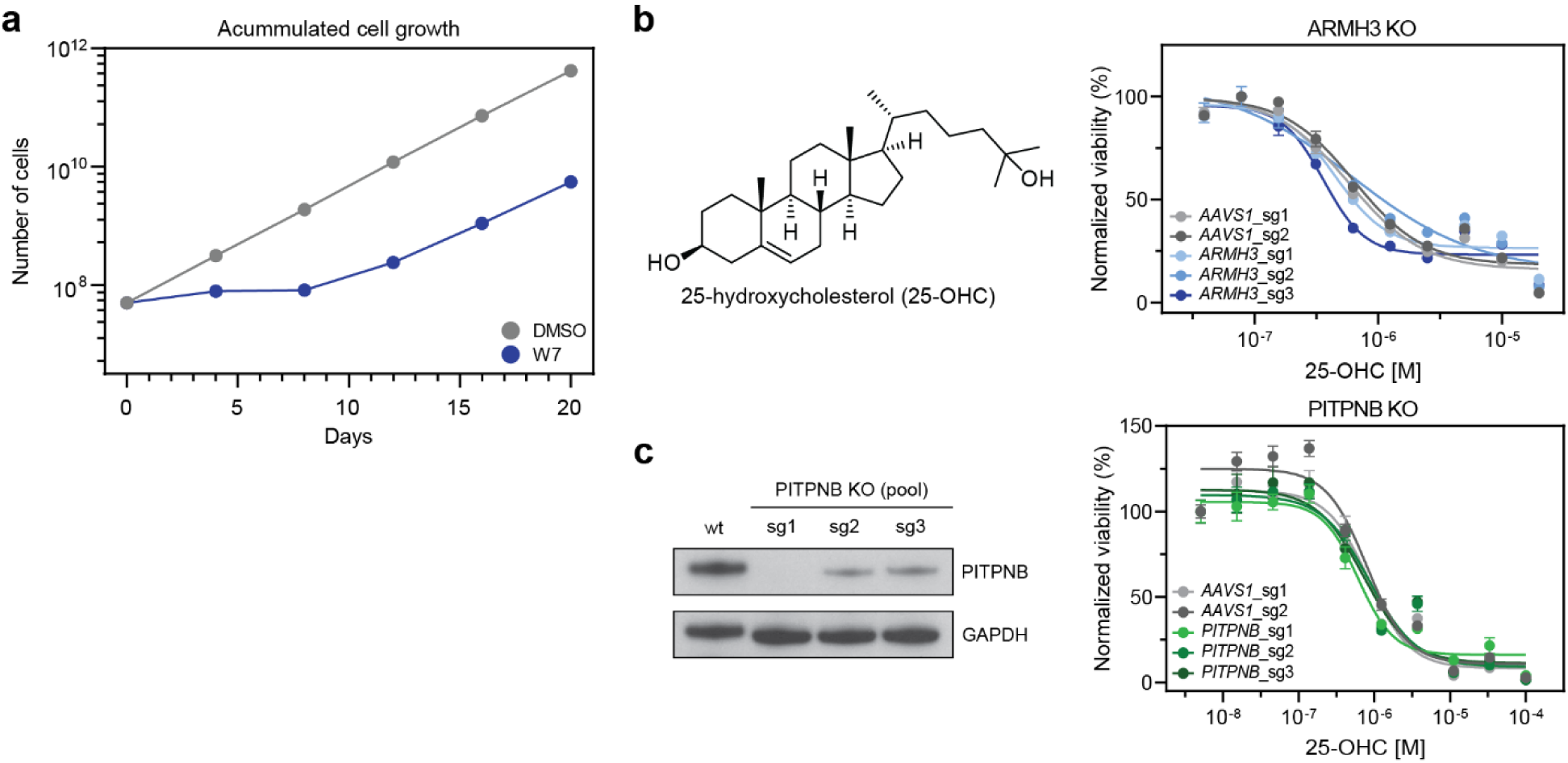
Genome-wide CRISPR/Cas9 resistance screen. **(a)** Accumulated cell growth curve of KBM7-Cas9 treated with DMSO or W7 over 20 days. **(b)** Dose-resolved, normalized viability after 72 h treatment of KBM7-Cas9 cells expressing sgRNAs targeting *ARMH3* (up, in blue), *PITPNB* (down, in green) or *AAVS1* locus (grey) with 25-OHC. Mean ± s.e.m.; n = 3 independent treatments. **(c)** PITPNB protein levels in KBM7 pools expressing three different sgRNAs against the gene. Representative images of n = 2 independent measurements.

**Extended Data Fig. 5.**
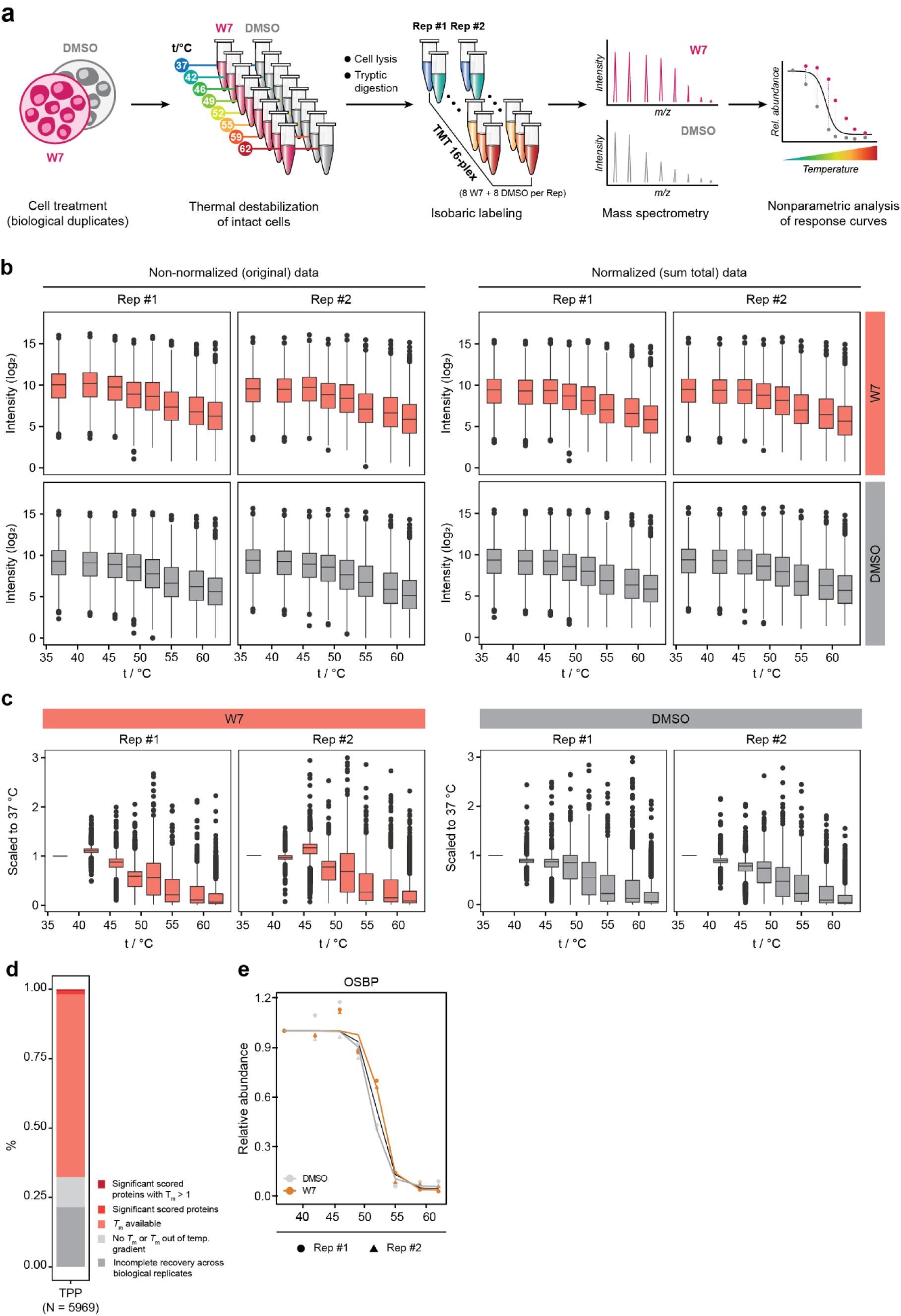
Thermal proteome profiling. **(a)** Overview of the experimental setup. Drug/vehicle treatment and 8-temperature destabilization were performed in biological duplicates. Significant shifts in drug-induced melting profiles were identified with NPARC. **(b)** Non-normalized (left) and sum total normalized (right) distribution of TPP data for individual W7/DMSO-treated replicates. **(c)** Distribution of TPP data after scaling each protein to the abundance signal in the lowest temperature condition (37 °C), shown for individual W7/DMSO-treated replicates. **(d)** Overview of TPP results represented as fraction of proteins in respect to individual analysis steps. **(e)** Melting profiles for the W7 hit OSBP detected by NPARC (adj. *P*-value < 0.01).

**Extended Data Fig. 6.**
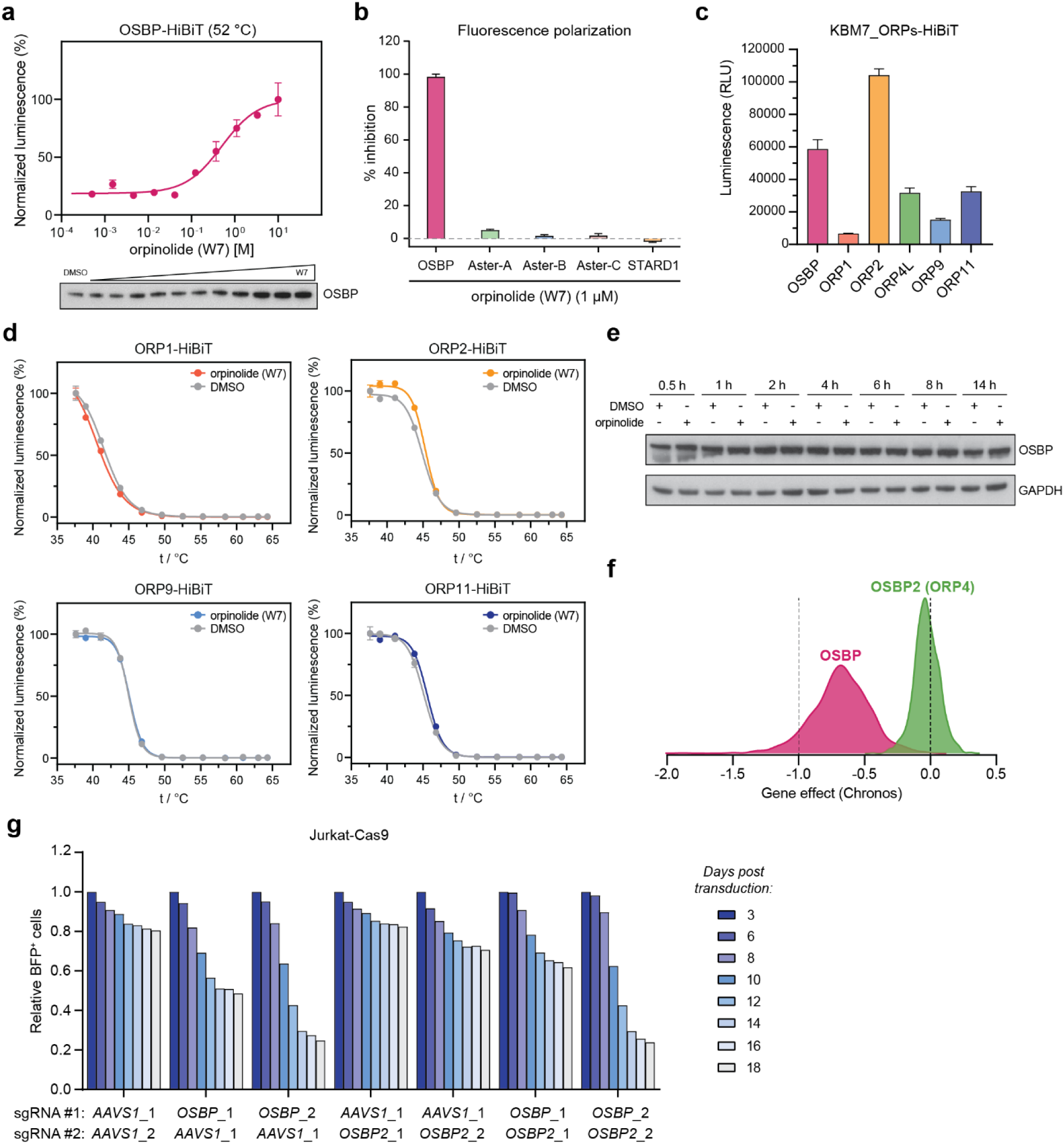
Oxysterol-binding protein OSBP is a putative target of orpinolide (W7) **(a)** Top: Dose-dependent stabilization of HiBiT-tagged OSBP at 52 °C in KBM7 cells stably expressing the fusion. Intact cells were treated for 4 h (highest conc. 10 μM, 1:3 drug dilutions) before performing the CETSA. Mean ± s.e.m.; n = 3 technical replicates; representative of n = 2 independent experiments is shown. Bottom: Endogenous OSBP levels in KBM7 after dose-resolved orpinolide treatment and thermal destabilization at 52 °C. Representative image of n = 2 independent measurements. **(b)** Fluorescence polarization competition assay measuring the displacement of 22-NDB-cholesterol from purified GST-tagged ORD of OSBP, Aster-A/B/C and STARD1 upon orpinolide treatment (1 μM). **(c)** Expression levels of different HiBiT-tagged ORPs in KBM7 cells expressing the fusions. Mean ± s.e.m.; n = 3 technical replicates. **(d)** Orpinolide does not thermally stabilize HiBiT-tagged ORP1, ORP2, ORP9 or ORP11 in KBM7 cells. Intact cells were treated for 4 h with 1 μM orpinolide before performing the CETSA. Mean ± s.e.m.; n = 3 technical replicates; representative of n = 2 independent experiments is shown. **(e)** Effect of orpinolide treatment (1 μM) on OSBP protein levels in KBM7 cells at different time points. Representative image of n = 2 independent measurements. **(f)** Distribution of OSBP and ORP4 (OSBP2) deletion effects across 1078 cancer cell lines from the DepMap 22Q4 CRISPR data. **(g)** CRISPR-based dropout screen in Jurkat cells constitutively expressing Cas9, transduced with a BFP-based dual knockout reporter. The reporter plasmid carries a combination of sgRNAs targeting *OSBP*, *OSBP2* or *AAVS1* locus to yield single or dual OSBP/ORP4 knockout. The bar plots depict relative BFP_+_ cells over 18 days, normalized to day 3 post transduction for each individual reporter system.

**Extended Data Table 1.**
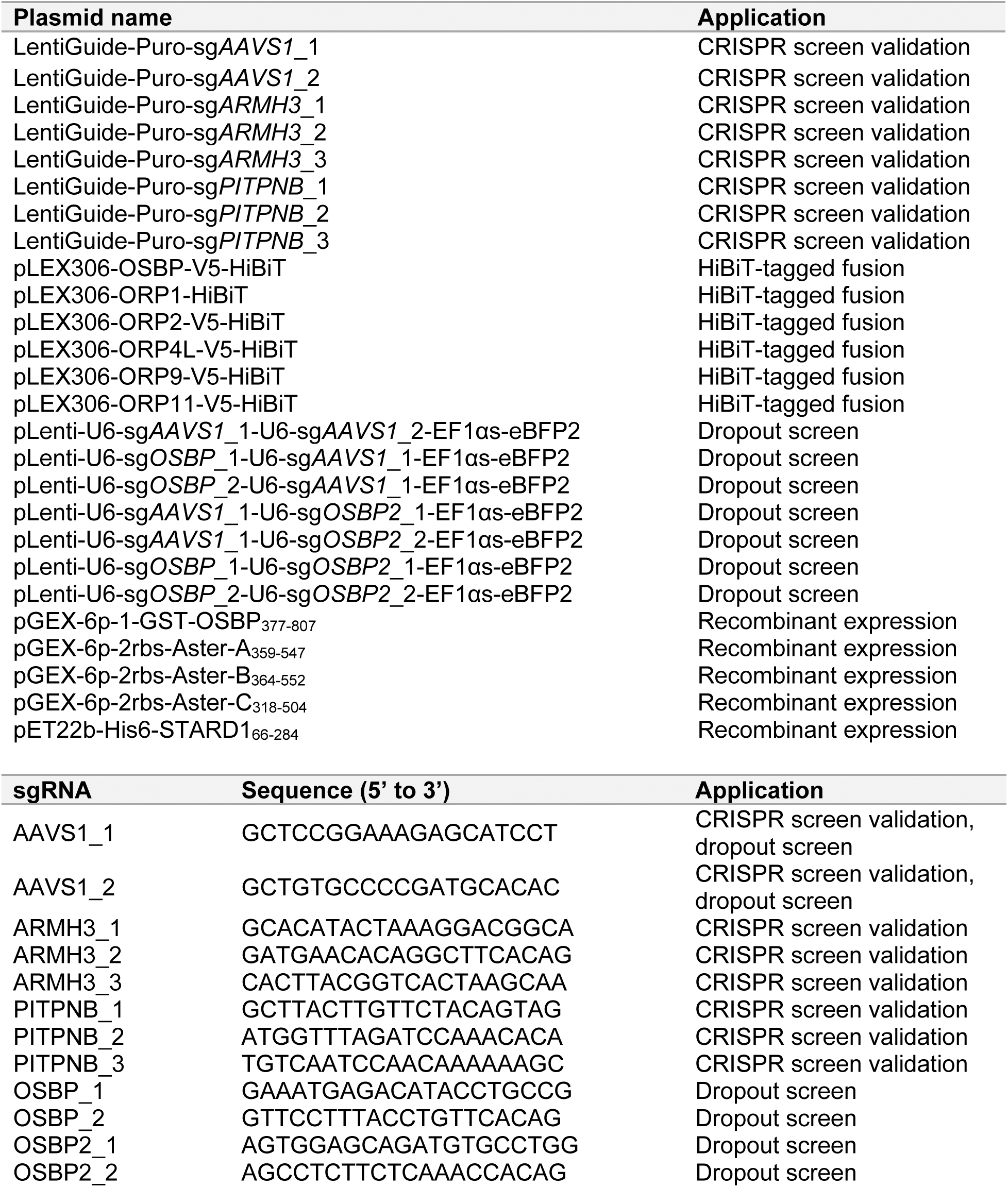
Plasmids and sgRNAs used in this study.

**Extended Data Table 2.**
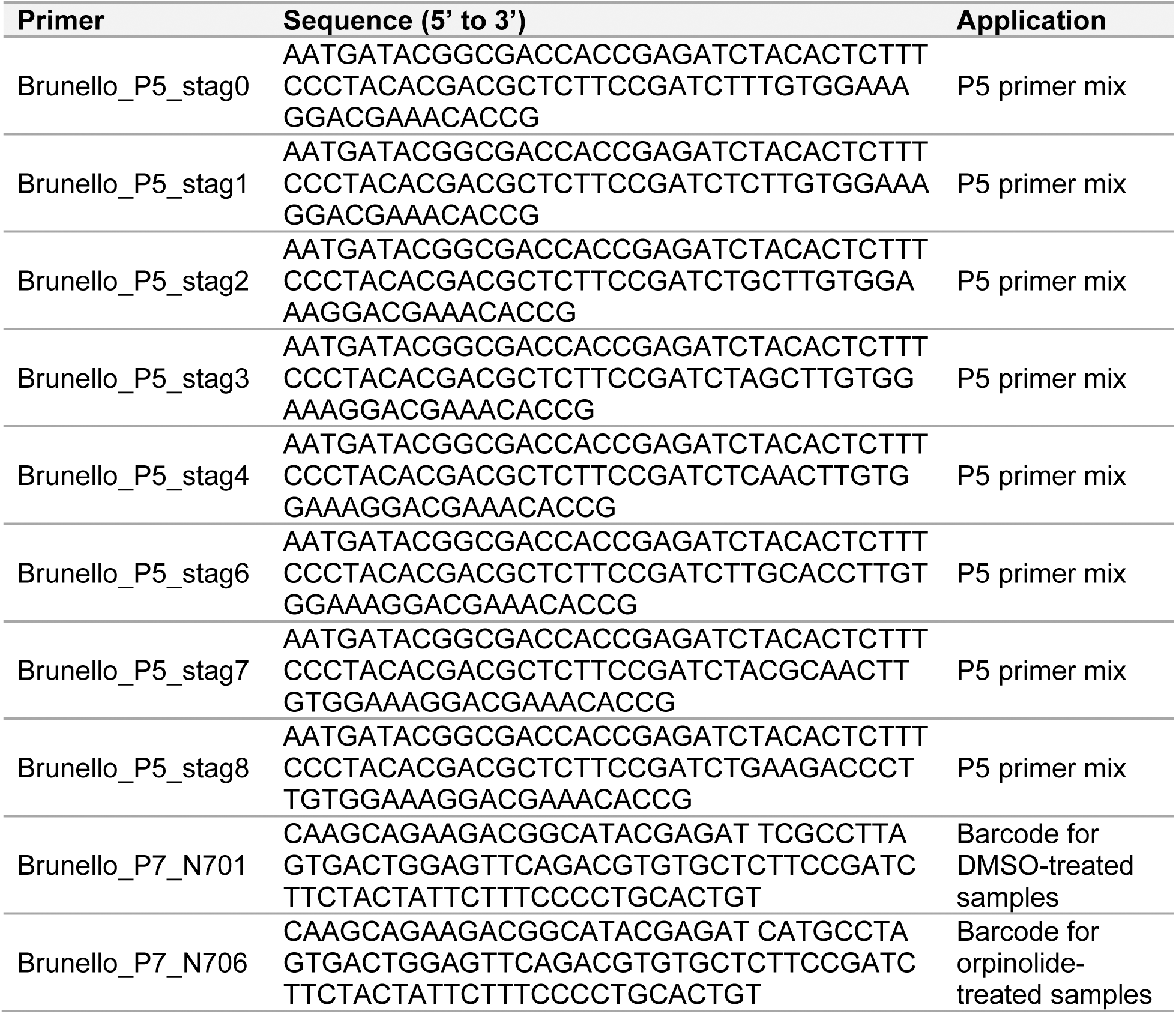
Primers used for PCR-amplification of sgRNAs in the genome-wide CRISPR/Cas9 screen.

## Methods

### Cell lines and cell culture

KBM7 cells were grown in IMDM supplemented with 10% FBS and 1% penicillin/streptomycin. K562, MV4;11, OCIAML3, NALM6, Jurkat, LOUCY, MOLT4 and P12-Ichikawa were grown in RPMI supplemented with 10% FBS and 1% Pen/Strep. Lenti-X 293T lentiviral packaging cells were grown in DMEM supplemented with 10% FBS and 1% Pen/Strep. KBM7 and Jurkat cells constitutively expressing Cas9 were generated as previously described.^51^ KBM7 iCas9 cells were a gift from Johannes Zuber. Non-malignant PBMCs were isolated from peripheral blood of a healthy adult volunteer (purchased from the local transfusion service, Red Cross Austria) via density gradient centrifugation (Lymphoprep, Stemcell Technologies). Cell lines were grown in a humidified incubator at 37 °C and 5% CO2 and were regularly tested for mycoplasma contamination.

### Plasmids, cloning and protein expression constructs

OSBP, ORP2, ORP4L, ORP9 and ORP11 cDNAs were custom synthesized by Twist Biosciences to include a C-terminal V5 tag and attB sites and cloned into pDONR221 (Thermo Fisher, 12536017). ORP1 cDNA was purchased in a Gateway compatible pGenDONR vector from GenScript. A pLEX306-HiBiT destination vector, kind gift from Michael Erb, was used for lentiviral expression of C-terminally tagged HiBiT fusions. *ARMH3*, *PITPNB*, *OSBP* and *OSBP2*-targeting sgRNAs were designed using the Vienna Bioactivity CRISPR score portal^52^ and were cloned either into LentiGuide-Puro (Addgene #52963) or into pLenti-U6-sgRNA#1-U6-sgRNA#2-EF1αs-eBFP2 (kind gift from Johannes Zuber) following standard protocol.^53, 54^ Human ASTER domains of Aster-A359-547, -B364-552 and -C318-504 were subcloned into a pGEX-6p-2rbs vector, thus introducing the cloning artifact ‘GPLGS’.^37^ The pGEX-6p-1-GST-OSBP377-807 plasmid was obtained from Genscript. The pET22b-His6-STARD166-284 plasmid was a gift from James H. Hurley (University of California).^55^ All generated plasmids and sgRNA sequences are listed in the Extended Data Table 1. The sequences of ORP cDNAs and synthesized fragments are listed in the Suppl. Table 6.

### Lentivirus production and transduction

Lenti-X 293T cells (at approx. 80% confluency) were co-transfected with the target vector, lentiviral psPAX2 helper (Addgene #12260) and pMD2.G envelope (Addgene #12259) using polyethyleneimine (PEI MAX^®^ MW 40,000, Polysciences, 24765-100) as previously described.^56^ Viral supernatant was harvested after 72 h and cleared of cellular debris by filtration through a 0.45-µm PES filter. Cells were transduced with respective virus via spinfection (2,000 rpm, 1 h, 37 °C) in the presence of polybrene (8 μg/mL, SantaCruz, SC-134220). For generation of ARMH3 and PITPNB knockouts in KBM7, cells were selected with puromycin (1 µg/mL, Gibco, A1113803) two days post transduction for a total of 5 days, after which the knockout pools were validated via western blotting.

### Phenotypic screening of the withanolide-inspired compound collection

The 52-membered withanolide-inspired compound collection (see Extended Data Fig. 1a and Suppl. Table 1) was designed and synthesized as previously reported.^15^ The compounds were printed onto 384-well plates (Corning 3570) in 10 concentration points (10, 3, 1, 0.3, 0.1, 0.03, 0.01 and 0.003 μM) and in biological triplicates. DMSO was used as a negative control, while staurosporine (1 μM) was used as a cytotoxic positive control. Both controls were seeded in multiple wells and in all of the screened plates.

Wild-type cells (K562, KBM7, MV4;11, OCIAML3, NAML6, Jurkat, LOUCY, MOLT4 and P12-Ichikawa) were seeded at the density of 1000 cells per well (50 μL assay volume) with the Liquid Handler Multidrop Combi. Cell viability was assessed after 72 h with the CellTiter-Glo assay (Promega, G7573). Survival curves and area under the curve (A.U.C.) were determined in GraphPad Prism (v9.5.1) by interpolation of a sigmoidal standard curve. Each data point was normalized to the mean luminescence of DMSO controls.

### Cell viability assay

Cells were seeded in 96-well plates at the density of 5000 cells per well and treated with DMSO or drug (1:3 serial dilution; ten different concentrations) for 72 h in biological triplicates. Cell viability was assessed with the CellTiter-Glo assay (Promega, G7573) according to manufacturer’s protocol. Luminescence signal was measured on the Multilabel Plate Reader Platform Victor X3 model 2030 (Perkin Elmer). Survival curves and half-maximum effective concentrations (EC50) were determined in GraphPad Prism (v9.5.1) by fitting a non-linear regression curve or interpolation of a sigmoidal standard curve. Each data point was normalized to the mean luminescence of DMSO controls or the lowest drug concentration.

### Cell painting assay

The cell painting assay was performed in a 384-well plate format in U2OS cells, in triplicates. Plate preparation, image acquisition, subsequent cellular feature extraction (579 in total) and compilation of phenotypic profiles were performed as previously described.^57^ The phenotypic profile of a compound is determined as the list of Z-scores of all features (calculated relative to the median of DMSO controls and defined by how many median absolute deviations the measured value is away from the median of the controls) for one compound. Additionally, an induction value was determined for each compound as the fraction of significantly changed features, in percent:

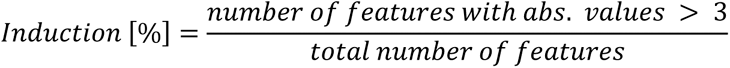

Similarities of phenotypic profiles (or Biosimilarity) were calculated from the correlation distances between two profiles (*Biosimilarity* = 1 – *Correlation Distance*; https://docs.scipy.org/doc/scipy/reference/generated/scipy.spatial.distance.correlation.html) and the compounds with the most similar profiles were determined from a set of 3000 reference compounds that was also measured in the assay. UMAP analysis was carried out as previously reported.^58^

### Western blotting

Collected cell pellets were washed with PBS and lysed in RIPA buffer (50 mM Tris-HCl pH 8.0, 150 mM NaCl, 1% Triton X-100, 0.5% sodium deoxycholate, 0.1% SDS, 1× Halt protease inhibitor cocktail, 25 U/mL Benzonase) by incubating on ice for at least 15 min. The lysates were cleared through centrifugation (15 min, 20,000g, 4 °C) and the total protein concentration determined by BCA protein assay (Pierce BCA Protein Assay Kit, Thermo Scientific, 23225) following manufacturer’s protocol. All lysates were supplemented with 4x Bolt LDS sample buffer (Invitrogen) and denatured for 5 min at 95 °C before loading onto polyacrylamide gels (20 μg of total protein). Proteins were separated on 4-12% SDS-PAGE gels (Invitrogen) and transferred to nitrocellulose membranes. The transfer efficiency was tested through staining with Ponceau-S. The membranes were blocked with 5% milk in TBST (30 min, RT). Primary antibodies were incubated overnight at 4 °C in 1% milk in TBST while the respective secondary antibodies were incubated in TBST for 1 h at RT. Blots were developed with chemiluminescence films. Following primary antibodies were used: OSBP (1:2,000; Bethyl, A304-553A), GAPDH (1:5,000; Santa Cruz Biotechnology, sc-365062), GOLIM4 (1:1,000; Thermo Scientific, PA5-51624), PITPNB (1:2,000; Bethyl, A305-591A). Secondary antibody: Peroxidase-conjugated AffiniPure Goat Anti-Rabbit IgG (1:10,000; Jackson ImmunoResearch 111-035-003).

### Expression proteomics

#### Sample preparation

KBM7 wt cells (40 million) were treated with orpinolide (1 μM) or DMSO for 8 h in biological duplicates. Afterwards, cells were collected via centrifugation, and the cell pellets washed four times with ice-cold PBS before being flash-frozen in liquid nitrogen. Each sample was lysed in 500 μL of lysis buffer (50 mM HEPES pH 8.0, 2% SDS, 1 mM PMSF, 1x protease inhibitor cocktail (Sigma-Aldrich)) for 20 mins at RT before heating to 99 °C for 5 min. After cooling down to RT, the DNA was sheared by sonication (Covaris S2 high performance ultrasonicator) and the lysates were cleared through centrifugation (20 min, 16,000g, 20 °C). Total protein concentration determined by BCA protein assay (Pierce BCA Protein Assay Kit, Thermo Scientific). Filter-aided sample preparation (FASP) was performed using a 30 kDa molecular weight cut-off filter (VIVACON 500; Sartorius Stedim Biotech). In brief, 200 μg total protein per sample were reduced with DTT (83.3 mM final concentration) followed by 5 min incubation at 99 °C. After cooling to RT, samples were mixed with 200 μL of freshly prepared UA solution (8 M urea in 100 mM Tris-HCl pH 8.5) and the filter units were centrifuged (15 min, 14,000g, 20 °C) to remove SDS. Residual SDS was washed out by a second washing step with 200 μL of UA. The proteins were alkylated with 100 μL of 50 mM iodoacetamide for 30 min at RT in the absence of light. Afterwards, three washing steps with 100 μL of UA solution were performed, followed by three washing steps with 100 μL of 50 mM TEAB buffer. Proteins were digested with trypsin at a ratio of 1:50 overnight at 37 °C. Peptides were recovered using 40 μL of 50 mM TEAB buffer, followed by 50 μL of 0.5 M NaCl, and were desalted using C18 solid phase extraction spin columns (The Nest Group). After desalting, peptides were labeled with TMT 16-plex reagents according to manufacturer’s protocol (Thermo Scientific Pierce). After quenching, all samples were pooled, organic solvent concentrated under vacuum and labeled peptides cleaned via C18 solid phase extraction (SPE).

#### Offline fractionation via RP-HPLC at high pH

Tryptic peptides were rebuffered in 10 mM ammonium formate (pH 10) prior to separation by reversed-phase (RP) liquid chromatography as previously described.^59^ Peptides were separated into 96 time-based fractions on a Phenomenex C18 RP column (150 × 2.0 mm, Gemini-NX 3 µm C18 110 Å; Phenomenex) using a Dionex UltiMate 3000 RSLCnano system, fitted with a binary high-pressure gradient pump, delivering solvent at 50 µL/min. Acidified fractions were consolidated into 36 fractions via a previously described concatenated strategy.^60^ After removal of solvent in a vacuum concentrator, samples were reconstituted in 0.1% TFA prior to LC-MS/MS analysis.

#### Data acquisition

LC-MS/MS analysis was performed on an Orbitrap Fusion Lumos Tribrid mass spectrometer (Thermo Scientific) coupled to a Dionex UltiMate 3000 RSLCnano system (Thermo Scientific) via a Nanospray Flex Ion Source (Thermo Scientific) interface. Peptides were loaded onto a trap column (PepMap 100 C18, 5 μm, 5 × 0.3 mm, Thermo Scientific) at a flow rate of 10 μL/min using 0.1% TFA as loading buffer. After loading, the trap column was switched in-line with a 75 µm inner diameter (ID) x 400 mm analytical column (packed in-house with ReproSil-Pur 120 C18-AQ, 3 µm particle size, Dr. Maisch HPLC GmbH). Mobile phase A consisted of 0.4% formic acid in water, while mobile phase B consisted of 0.4% formic acid in a mixture of 90% acetonitrile and 9.6% water. Separation was achieved using a multistep gradient over 90 min at a flow rate of 230 nL/min (increase of initial gradient from 6% to 9% solvent B within 1 min, 9% to 30 % solvent B within 81 min, 30% to 65% solvent B within 8 min, 65% to 100% solvent B within of 1 min and 100% solvent B for 6 mins before equilibrating to 6% solvent B for 18 mins prior to next injection). In the liquid junction setup, electrospray ionization was enabled by applying a voltage of 1.8 kV directly to the liquid being sprayed, and non-coated silica emitter was used. The mass spectrometer was operated in a data-dependent acquisition mode (DDA) and used a synchronous precursor selection (SPS) approach. For both MS2 and MS3 levels, we collected a 400–1600 m/z survey scan in the Orbitrap at 120,000 resolution (FTMS1), the AGC target was set to “standard” and a maximum injection time (IT) of 50 ms was applied. Precursor ions were filtered by charge state (2-5), dynamic exclusion (120 s with a ±10 ppm window), and monoisotopic precursor selection. Precursor ions for data-dependent MSn (ddMSn) analysis were selected using 10 dependent scans (TopN approach). A charge-state filter was used to select precursors for data-dependent scanning. In ddMS2 analysis, spectra were obtained using one charge state per branch (from z=2 to z=5) in a dual-pressure linear ion trap (ITMS2). The quadrupole isolation window was set to 0.7 Da and the collision-induced dissociation (CID) fragmentation technique was used at a normalized collision energy of 34%. The normalized AGC target was set to standard with a maximum IT of 35 ms. During the ddMS3 analyses, precursors were isolated using SPS waveform and different MS1 isolation windows (1.3 m/z for z=2, 1.2 m/z for z=3, 0.8 m/z for z=4 and 0.7 m/z for z=5). Target MS2 fragment ions were further fragmented by high-energy collision induced dissociation (HCD) followed by orbitrap analysis (FTMS3). The normalized HCD collision energy was set to 45% and the normalized AGC target was set to 300% with a maximum IT set to “auto”. The resolution was set to 50,000 with a defined scanning range of 100 to 500 m/z. Xcalibur (v3.3.2782.34) and Tune (v3.3) were used to operate the instrument.

#### Data analysis

Acquired raw data files were processed with the Proteome Discoverer (v2.4.1.15) utilizing the Sequest HT database search engine and Percolator validation software node to remove false positives with a false discovery rate (FDR) of 1% on peptide and protein level under strict conditions. Searches were performed with full tryptic digestion against the human SwissProt database (Homo sapiens (SwissProt TaxID=9606) (v2017), 42252 sequences) with or without deamidation (+0.9840 Da) on amino acids Asp and Glu. Met oxidation (+15.994 Da) and protein N-terminal acetylation (+42.011 Da), as well as Met loss (−131.040 Da) and protein N-terminal acetylation with Met loss (−89.030 Da) were set as variable modifications, while carbamidomethylation (+57.021 Da) of Cys residues and TMT 16-plex labeling of peptide N-termini and Lys residues (+304.207 Da) were set as fixed modifications. Data was searched with mass tolerances of ± 10 ppm and ± 0.6 Da on the precursor and fragment ions, respectively. Results were filtered to include peptide spectrum matches (PSMs) with Sequest HT cross-correlation factor (Xcorr) scores of ≥ 1 and high peptide confidence assigned by Percolator. MS3 signal-to-noise (S/N) values of TMT reporter ions were used to estimate peptide/protein abundance changes. PSMs with precursor isolation interference values of ≥ 70%, SPS mass matches ≤ 65%, and average TMT reporter ion S/N ≤ 10 were excluded from quantitation. Both unique and razor peptides were used for TMT quantitation. Isotopic impurity correction and TMT channel-normalization based on total peptide amount were applied. For statistical analysis and P-value calculation, the integrated ANOVA hypothesis test was used. TMT ratios with P-values below 0.05 were considered as significant. Only proteins with > 1 peptide detected and > 1 protein unique peptide detected were considered for further analysis. Gene Ontology (GO) enrichment analysis of downregulated proteins was performed using tools available on the GO Consortium website (http://geneontology.org; release 2023-01-01).

### Protein interaction analysis of up- and downregulated proteins

A full proteome profiling, employing TMT technique, of orpinolide versus DMSO treated cells, identified 59 up- and 116 downregulated proteins (175 regulated proteins; log2 fold change greater than 0.25 or less than −0.25, adj. *P*-value < 0.05). Localization of each protein was annotated by combining the localization information form The Human Protein Atlas project (v22.0)^61^ and UniProtKB (13.01.2023). The different localization information was further manually curated, considering the multiple localizations and corresponding confidence levels. From the interaction database BioGRID (v4.4.212)^23^, a protein-protein interaction (PPI) network limited to direct physical interactions between the regulated proteins was derived. For this, first the BioGRID database was filtered, to only include interactions reported for homo sapiens (9606). Further, only interactions which were annotated to be generated by “Affinity Capture-Western”, “Co-purification” and “Affinity Capture-MS” were used, omitting all proximity interaction and co-elution profiling technologies. This resulted in an interaction database with 610,485 interactions. The filtered BioGRID database contained nodes for 172 of the 175 regulated proteins. Next, all edges between the regulated proteins were extracted resulting in a network covering 97 nodes (55% of all regulated proteins) and 97 edges connecting the nodes. 75 regulated proteins were not directly connected to other reported regulated proteins. We sampled 10,000 times 172 random nodes from the filtered BioGRID database and obtained a higher number of nodes (permutation *P*-value = 0.0097) for the reconstructed network of regulated proteins, while none of the permutated networks had a higher or equal number of edges (permutation *P*-value = 0), indicating that the up- and downregulated proteins represent a significantly more than random related set of proteins. Data analysis was conducted with the statistical software R (version R-4.2.0) while the network was visualized using the software Cytoscape (v.3.8.0).

### RNA sequencing

KBM7 wt cells (10 million) were treated with orpinolide (485 nM) or DMSO for 6 h in biological triplicates. Total RNA was extracted with the RNeasy^®^ Mini Kit (Qiagen, 74106) and its amount determined using Qubit RNA HS kit (Thermo, Q32852). Next, Poly(A) enrichment (Lexogen No. 039) was performed with 4 µg of RNA per condition and the RNA-seq library was prepped using the Corall Total RNA-Seq Library Prep Kit (Lexogen No. 095) and following the manufacturer’s protocol (2 ng RNA as starting material). The endpoint PCR was performed with 16 cycles. Amplified and purified sequencing libraries were analyzed on an Agilent 2100 Bioanalyzer (High sensitivity DNA analysis kit, Agilent, 5067-4626), pooled in equimolar amounts (2.47 ng/μL), and sequenced using the Illumina HiSeq 4000 platform and 50 bp single-end configuration.

#### RNA sequencing data analysis

Raw reads were trimmed using Trimmomatic (v0.32).^62^ The following parameters were used: HEADCROP:13 ILLUMINACLIP:epignome_adapters_2_add.fa:2:10:4:1:true SLIDINGWINDOW:4:1 MAXINFO:16:0.40 MINLEN:30. Trimmed reads were mapped to the hg38/GRCh38 assembly of the human reference genome using STAR aligner (v2.5.2b)^63^ using the following parameters: --outFilterType BySJout -- outFilterMultimapNmax 20 --alignSJoverhangMin 8 --alignSJDBoverhangMin 1 -- outFilterMismatchNmax 999 --outFilterMismatchNoverLmax 0.6 --alignIntronMin 20 -- alignIntronMax 1000000 --alignMatesGapMax 1000000 --outSAMattributes NH HI NM MD --outSAMtype BAM SortedByCoordinate. Genes were defined based on genome-build GRCh38.p7, genome-build-accession NCBI: GCA_000001405.22. Read counts per gene were obtained from the aligned reads using the htseq-count command (v0.11.2) from the HTSeq framework.^64^ The Bioconductor/R package DESeq2 (v1.34.0)^65^ was used for normalization and differential gene expression analysis. The Gene Sets Enrichment Analysis (GSEA) was performed by the GSEA software (v3.0)^66^ and the results represented using R. To build the gene rankings, genes were ranked based on meta-score values, the meta-score represents “the worst” score from the following three scores obtained from the DESeq2 analysis: (i) Log2FC, (ii) baseMean RNA-seq score, (iii) p-value. The following GSEA parameters were used: xtools.gsea.GseaPreranked -nperm 1000 -scoring_scheme weighted -norm meandiv.

### Genome-wide CRISPR/Cas9 knockout screen

Genome-wide CRISPR/Cas9 positive selection screens were performed as previously reported.^51^ KBM7 cells constitutively expressing Cas9 (KBM7-Cas9, 250 million) were transduced with the Brunello sgRNA library (Addgene #12260) at a multiplicity of infection (MOI) of 0.23 to yield a calculated library representation of 758 cells per sgRNA. Transductions were performed as described above. On the next day, the transduced cells were pooled and diluted with fresh IMDM. Pools were selected with puromycin (1 μg/mL) for 5 days, after which drug treatments were started. Selected pools (50 million cells; 0.5 million cells/mL density) were treated with DMSO or orpinolide (2x EC50 starting concentration). Every four days cells were counted, 50 million were reseeded (maintaining 0.5 million cells/mL density) and treated with fresh orpinolide or DMSO. Orpinolide concentration was dynamically adjusted to the accumulated growth curves to maintain a consistent impact on cell proliferation without losing the coverage. After 20 days of treatment, Lymphocyte Separation Medium (LSM, Corning) was used to remove dead cells and cellular debris. Viable cells were harvested and the cell pellets flash-frozen in liquid nitrogen.

#### Library preparation for next generation sequencing

Genomic DNA (gDNA) was isolated from respective samples with the QIAamp DNA Mini Kit (Qiagen, 51306) and used as template for PCR amplification of sgRNA sequences in batches of 20 μg of gDNA per PCR reaction. One PCR reaction (100 μL) furthermore contained P5 forward primer mix (0.5 μM), condition-specific P7 barcoded primer (0.5 μM), ExTaq polymerase (Clontech, 1.5 μL), dNTP mix (8 μL), 10x buffer (10 μL) and water. Used primer sequences are listed in the Extended Data Table 2. Target amplification: 1 min at 95 °C initial denaturation; 30 sec at 95 °C, 30 sec at 53 °C, 30 sec at 72 °C for 27 cycles; 10 min at 72 °C final elongation. After pooling all PCR reactions for specific condition, the 360-bp amplicon was purified with AMPure XP beads (Beckman Coulter, 10136224; 0.7x right side size selection) and eluted with TE buffer (28 μL). Sequencing libraries were analyzed on an Agilent 2100 Bioanalyzer (High sensitivity DNA analysis kit, Agilent, 5067-4626) pooled in equimolar amounts (2.8 ng/μL) and sequenced using the Illumina HiSeq 4000 platform and 50 bp single-end configuration.

#### Next-generation sequencing data analysis

Raw read files were converted to fastq format using the convert function from BamTools (v2.5.2).^67^ Sequencing adapters were trimmed using Cutadapt (v3.4)^68^ with -g CGAAACACCG and --minimum-length = 10. The 20 bp of spacer sequence were then extracted using fastx toolkit (v0.0.14) (http://hannonlab.cshl.edu/fastx_toolkit/) and aligned to the respective sgRNA index using Bowtie2 (v2.4.4)^69^ allowing for one mismatch in the seed sequence. Spacers were counted using the bash command ‘cut -f 3 (0) | sort | uniq -c’ on the sorted SAM files. A count table with all conditions was then assembled, and the counts + 1 were converted to counts-per-million to normalize for sequencing depth. Log2-normalized fold changes compared to DMSO were calculated for each spacer. Statistical analysis was performed using the STARS algorithm (v1.3).^27^ For this, spacers were rank-ordered based on log2 fold change and tested with the parameters -- thr 10 --dir P against a null hypothesis of 1000 random permutations. Genes with FDR < 0.05 were called as hits.

### Thermal proteome profiling (TPP) in intact cells

#### Sample preparation

KBM7 cells (36 million, 1.5 million cells/mL density) were treated with orpinolide (5 μM) or DMSO for 2 h in biological duplicates. Afterwards, cells were collected via centrifugation, washed three times with ice-cold PBS and transferred into PCR plates so that each well contained 4 million cells (100 μL). After spinning down, 80 μL of the supernatant were removed without disturbing the pellet. The samples were then thermally destabilized (3 min 8-temperature gradient (37.3, 42.7, 46.2, 49.4, 52.7, 55.9, 59.4 and 62.3 °C) followed by 3 min at 25 °C) and re-suspended in 60 uL of the lysis buffer (20 mM Tris-HCl pH 8.0, 120 mM NaCl, 0.5% NP-40, 1× Halt protease inhibitor cocktail). Four freeze/thaw cycles with liquid nitrogen were then performed and the lysed samples were cleared out via centrifugation (full speed, 1 h, 4 °C). In total, 38 lysates were obtained with each biological replicate comprising 16 DMSO/orpinolide-treated samples destabilized at 8 different temperatures. Cleared supernatants (70 μL) were supplemented with SDS to a final concentration of 2% and the protein concentration determined by BCA protein assay (Pierce BCA Protein Assay Kit, Thermo Scientific). Filter-aided sample preparation (FASP) was performed as described in the *Expression proteomics* section, using a 30 kDa molecular weight cut-off filter. After reduction and alkylation, proteins (in 50 μL of 50 mM TEAB buffer) were digested with trypsin (3 μg per sample; protein input for digest between 40-150 μg depending on temperature condition) overnight at 37 °C, followed by addition of fresh trypsin (4.5 μg per sample) for further three hours. Peptides were recovered using 40 μL of 50 mM TEAB buffer, followed by 50 μL of 0.5 M NaCl, and were desalted using Pierce Peptide Desalting Spin Columns (Thermo Scientific). Desalted peptides were subsequently labeled with TMT 16-plex reagents (two 16-plexes in total, one for each biological replicate) according to manufacturer’s protocol (Thermo Scientific Pierce). One TMT 16-plex comprised of 8 temperature points of orpinolide and 8 of DMSO treated cells. After quenching, all samples per respective 16-plex were pooled, organic solvent concentrated under vacuum and labeled peptides cleaned via C18 solid phase extraction (SPE).

#### Offline fractionation and data acquisition

Offline fractionation via RP-HPLC at high pH (20 fractions per 16-plex) was performed as described in the *Expression proteomics* section.

LC-MS/MS analysis was performed on an Orbitrap Fusion Lumos Tribrid mass spectrometer (Thermo Scientific) coupled to a Dionex UltiMate 3000 RSLCnano system (Thermo Scientific) via a Nanospray Flex Ion Source (Thermo Scientific) interface. Peptides were loaded onto a trap column (PepMap 100 C18, 5 μm, 5 × 0.3 mm, Thermo Scientific) at a flow rate of 10 μL/min using 0.1% TFA as loading buffer. After loading, the trap column was switched in-line with an Acclaim PepMap nanoHPLC C18 analytical column (2.0 µm particle size, 75 µm ID x 500 mm, catalog number 164942, Thermo Scientific). The column temperature was maintained at 50 °C. Mobile phase A consisted of 0.4% formic acid in water, while mobile phase B consisted of 0.4% formic acid in a mixture of 90% acetonitrile and 9.6% water. Separation was achieved using a multistep gradient over 90 min at a flow rate of 230 nL/min (increase of initial gradient from 6% to 9% solvent B within 1 min, 9% to 30% solvent B within 81 min, 30% to 65% solvent B within 8 min, 65% to 100% solvent B within of 1 min and 100% solvent B for 6 mins before equilibrating to 6% solvent B for 18 mins prior to next injection). In the liquid junction setup, electrospray ionization was enabled by applying a voltage of 1.8 kV directly to the liquid being sprayed, and non-coated silica emitter was used. The mass spectrometer was operated in a data-dependent acquisition mode (DDA) and used a synchronous precursor selection (SPS) approach. For both MS2 and MS3 levels, we collected a 400– 1650 m/z survey scan in the Orbitrap at 120,000 resolution (FTMS1), the AGC target was set to “standard” and a maximum injection time (IT) of 50 ms was applied. Precursor ions were filtered by charge state (2-6), dynamic exclusion (60 s with a ±10 ppm window), and monoisotopic precursor selection. Precursor ions for data-dependent MSn (ddMSn) analysis were selected using 10 dependent scans (TopN approach). A charge-state filter was used to select precursors for data-dependent scanning. In ddMS2 analysis, spectra were obtained using one charge state per branch (from z=2 to z=5) in a dual-pressure linear ion trap (ITMS2). The quadrupole isolation window was set to 0.7 Da and the collision-induced dissociation (CID) fragmentation technique was used at a normalized collision energy of 35%. The normalized AGC target was set to 200% with a maximum IT of 35 ms. During the ddMS3 analyses, precursors were isolated using SPS waveform and different MS1 isolation windows (1.3 m/z for z=2, 1.2 m/z for z=3, 0.8 m/z for z=4 and 0.7 m/z for z=5). Target MS2 fragment ions were further fragmented by high-energy collision induced dissociation (HCD) followed by orbitrap analysis (FTMS3). The normalized HCD collision energy was set to 45% and the normalized AGC target was set to 300% with a maximum IT set to “auto”. The resolution was set to 50,000 with a defined scanning range of 100 to 500 m/z. Xcalibur (v4.3.73.11) and Tune (v3.4.3072.18) were used to operate the instrument.

#### Data analysis

First, common lab contaminants were filtered from the datasets and the abundance values of the biological replicates. Next, we normalized all measurements (n = 4) per temperature point by applying a sum total normalization (see Extended Data Fig. 4b). Only proteins quantified ≥ 2 PSMs across the DMSO (vehicle) and compound treated conditions were considered (removal of 470 ProteinGroups across all conditions). Next, the relative abundance of each protein per condition and replicate was scaled to 37 °C, the lowest temperature point. Subsequently, we employed the nonparametric analysis of response curves (NPARC) R-package^34^ to identify significant shifts in protein melting behavior upon compound treatment. The NPARC method has the advantage to identify subtle but highly reproducible melting temperature (*T*m) shifts by comparing protein melting curves, including goodness-of-fit and residual errors of the model, and does not rely on the estimated *T*m point. Prior to model fitting, the dataset was filtered to proteins quantified in both biological replicates. This resulted in 4,682 proteins (from 5,969 in total) which were used for the NPARC method. We obtained *in vivo* melting curves for 4,037 proteins, whereas 113 were found to show a significant shift in *T*m with an adjusted *P*-value below 0.01. The obtained hits were further filtered to yield proteins with a Δ*T*m exceeding 1 °C (see Suppl. Table 5).

### Cellular thermal shift assay (CETSA) in intact cells

After treatment with orpinolide or DMSO (1 million cells/mL), KBM7 wt or KBM7 overexpressing HiBiT-tagged ORPs were spun down (1450 rpm, 5 min), washed once with PBS, and re-suspended in PBS (100 uL of PBS per 1 million of cells). The cell suspension (100 μL) was distributed into PCR strips, spun down (1450 rpm, 5 min) and 80 μL of the supernatant were removed without disturbing the cell pellet. The samples were then thermally destabilized (3-min temperature gradient of choice followed by 3 min at 25 °C) and re-suspended in 30 uL of the lysis buffer (20 mM Tris-HCl pH 8.0, 120 mM NaCl, 0.5% NP-40). Three freeze/thaw cycles with liquid nitrogen were then performed and the lysed samples were cleared out via centrifugation (full speed, 20 min, 4 °C).

#### Analysis via immunoblotting

Cleared supernatants were supplemented with 4x Bolt LDS sample buffer (Invitrogen) and denatured for 5 min at 95 °C before loading (6 μL) onto 4-12% polyacrylamide gels. Western blotting was performed as described above.

#### HiBiT-CETSA

Luminescence was assessed with the Nano-Glo^®^ HiBiT Lytic Detection System (Promega, N3040) in 384-well plate format and in technical triplicates. Briefly, a 6x detection master mix was prepared containing the substrate, LgBiT and the provided lysis buffer and 2.5 μL were distributed into well plates. Cleared lysates of cells expressing HiBiT-tagged ORPs (12.5 μL) were then added into respective wells and the plates were incubated at room temperature for 10 mins with mild shaking. Luminescence signal was measured on the Multilabel Plate Reader Platform Victor X3 model 2030 (Perkin Elmer). Melting curves and *T*m were determined in GraphPad Prism (v9.5.1) by fitting a non-linear regression curve or interpolation of a sigmoidal standard curve. Each data point was normalized to the mean luminescence at the lowest temperature (typically 37.6 °C).

### Recombinant proteins

#### Protein expression and purification

ASTER domains of human Aster-A359-547, -B364-552 and -C318-504 in the pGEX-6p-2rps vector with an N-terminal PreScission-cleavable GST-tag were expressed in *Escherichia coli* OverExpress C41 in Terrific Broth (TB) medium for approximately 16 h at 18 °C after induction with 0.1 mM IPTG. Cells were collected at 3,500g for 15 min and lysed by sonication in buffer containing 50 mM HEPES pH 7.5, 300 mM NaCl, 10% (vol/vol) glycerol, 5 mM DTT, 0.1% (vol/vol) Triton X-100 and protease inhibitor mix HP plus (Serva). The cleared lysate was purified by affinity chromatography on a GSTrap FF column (Cytiva) using an ÄKTA Start (Cytiva) in buffer containing 50 mM HEPES pH 7.5, 300 mM NaCl, 10% (vol/vol) glycerol, 5 mM DTT and 0.01% (vol/vol) Triton X-100. The GST-tag was cleaved on the column overnight at 4 °C. Proteins were further purified by size-exclusion chromatography on a HiLoad 16/600 Superdex 75 pg (Cytiva) in buffer containing 20 mM HEPES pH 7.5, 300 mM NaCl, 10% (vol/vol) glycerol and 2 mM DTT.

The START domain of human STARD166-284 harboring an N-terminal His6-Tag was expressed in *E. coli* BL21(DE3) in Luria-Bertani Broth (LB) medium for approximately 16 h at 18 °C after induction with 0.15 mM IPTG. Cells were collected at 3,500g for 15 min and lysed by sonication in buffer containing 50 mM HEPES pH 7.5, 150 mM NaCl, 5% (vol/vol) glycerol, 5 mM DTT, 0.1% (vol/vol) Triton X-100 and EDTA-free protease inhibitor cocktail (Sigma-Aldrich). The cleared lysate was purified by affinity chromatography on a Ni-NTA Superflow Cartridge (Qiagen) using an ÄKTA Start (Cytiva) in buffer containing 50 mM HEPES pH 7.5, 150 mM NaCl, 5% (vol/vol) glycerol, 5 mM DTT. STARD166-284 was eluted with buffer containing 50 mM HEPES pH 7.5, 150 mM NaCl, 5% (vol/vol) glycerol, 5 mM DTT and 500 mM imidazole. Proteins were further purified by size-exclusion chromatography on a HiLoad 16/600 Superdex 75 pg (Cytiva) in buffer containing 20 mM HEPES pH 7.5, 150 mM NaCl, 5% (vol/vol) glycerol and 2 mM DTT.

The ORP domain of human OSBP377-807 in the pGEX-6p-1 vector with an N-terminal PreScission-cleavable GST-tag was expressed in *E. coli* OverExpress C41 in LB medium for approximately 16 h at 18 °C after induction with 0.1 mM IPTG. Cells were collected at 3,500g for 15 min and lysed by sonication in buffer containing 20 mM HEPES pH 7.5, 300 mM NaCl, 10% (vol/vol) glycerol, 5 mM DTT, 0.1% (vol/vol) Triton X-100 and EDTA-free protease inhibitor cocktail (Sigma-Aldrich). The cleared lysate was purified by affinity chromatography on a GSTrap HF column (Cytiva) using an ÄKTA Start (Cytiva) in buffer containing 20 mM HEPES pH 7.5, 300 mM NaCl, 10% (vol/vol) glycerol, 5 mM DTT. OSBP377-807 was eluted with buffer containing 20 mM HEPES pH 7.5, 300 mM NaCl, 10% (vol/vol) glycerol, 5 mM DTT and 10 mM reduced glutathione. Proteins were further purified by size-exclusion chromatography on a HiLoad 16/600 Superdex 75 pg (Cytiva) using an ÄKTA Explorer (Cytiva) in buffer containing 20 mM HEPES pH 7.5, 150 mM NaCl, 10% (vol/vol) glycerol and 2 mM DTT.

### Fluorescence polarization

Fluorescence polarization (FP) experiments were performed in a buffer composed of 20 mM HEPES pH 7.5, 300 mM NaCl, 0.01% (vol/vol) Tween-20, 0.5% glycerol and 2 mM DTT in a final volume of 30 µL in black, flat-bottom, non-binding 384-well plates (Corning). For competition experiments, 20 nM 22-NBD-cholesterol was mixed with protein and incubated with desired concentrations of screening compounds. The fluorescence polarization signal was measured using a Spark Cyto multimode microplate reader (Tecan) with filters set at 485 ± 20 nm for excitation and at 535 ± 20 nm for emission. The data was analyzed using GraphPad Prism (v.9.5.1). Measured mP values were normalized setting 100% inhibition as the FP signal from the protein + fluorophore control well and 0% as the FP signal from the fluorophore control well. Curves were fitted to the normalized data via non-linear regression to allow the determination of IC50 values.

### Flow cytometry analysis of OSBP/ORP4 essentiality

To quantify the influence of OSBP and/or ORP4 genetic perturbation, KBM7 cells expressing inducible Cas9 (iCas9) or Jurkat cells constitutively expressing Cas9 were transduced with pLenti-U6-sgRNA#1-U6-sgRNA#2-EF1αs-eBFP2 reporter with 50-60% infection efficiency. The reporter plasmid carries a combination of sgRNAs targeting *OSBP*, *OSBP2* or *AAVS1* locus to yield single or dual OSBP/OSBP2 knockouts (see Extended Data Table 1). In KBM7 iCas9 cells, infection levels were determined by flow cytometry three days after transduction based on BFP marker expression (Day 0) and Cas9 expression was induced with doxycycline (0.4 µg/mL). In Jurkat-Cas9 cells, infection levels were determined by flow cytometry three days after transduction. In both cases, the percentage of sgRNA^+^ (BFP^+^) cells were monitored by flow cytometry in regular intervals. Flow cytometry measurements were performed on LSRFortessa (BD Biosciences) while the data was analyzed in FlowJo (v10.8.1).

### Data availability

The mass spectrometry proteomics data (Figures 1g-1h and 3a, Extended Data Fig. 3 and 5, Suppl. Tables 2 and 5) have been deposited to the ProteomeXchange Consortium via the PRIDE^70^ partner repository with the dataset identifier PXD040694 and 10.6019/PXD040694 (expression proteomics) as well as PXD040692 and 10.6019/PXD040692 (thermal proteome profiling). Raw and analyzed RNA-seq and genome-wide CRISPR/Cas9 screening datasets (Figure 2, Extended Data Fig. 4, Suppl. Tables 3 and 4) are available in NCBI’s Gene Expression Omnibus under accession number GSE226849. The data supporting all of the findings in this study are available within the paper, its supplementary files, and the mentioned databases. The data analysis codes are available at https://github.com/GWinterLab/W7.

